# Inferring the Demographic History of Inbred Species From Genome-Wide SNP Frequency Data

**DOI:** 10.1101/2019.12.20.881474

**Authors:** Paul D. Blischak, Michael S. Barker, Ryan N. Gutenkunst

## Abstract

Demographic inference using the site frequency spectrum (SFS) is a common way to understand historical events affecting genetic variation. However, most methods for estimating demography from the SFS assume random mating within populations, precluding these types of analyses in inbred populations. To address this issue, we developed a model for the expected SFS that includes inbreeding by parameterizing individual genotypes using beta-binomial distributions. We then take the convolution of these genotype probabilities to calculate the expected frequency of biallelic variants in the population. Using simulations, we evaluated the model’s ability to co-estimate demography and inbreeding using one- and two-population models across a range of inbreeding levels. We also applied our method to two empirical examples, American pumas (*Puma concolor*) and domesticated cabbage (*Brassica oleracea* var. *capitata*), inferring models both with and without inbreeding to compare parameter estimates and model fit. Our simulations showed that we are able to accurately co-estimate demographic parameters and inbreeding even for highly inbred populations (*F* = 0.9). In contrast, failing to include inbreeding generally resulted in inaccurate parameter estimates in simulated data and led to poor model fit in our empirical analyses. These results show that inbreeding can have a strong effect on demographic inference, a pattern that was especially noticeable for parameters involving changes in population size. Given the importance of these estimates for informing practices in conservation, agriculture, and elsewhere, our method provides an important advancement for accurately estimating the demographic histories of these species.

## Introduction

Estimating the demographic history of closely related populations or species is an important first step in understanding the interplay of the evolutionary forces shaping genetic variation. Divergence, migration, changes in population size, and other historical events all contribute to population allele frequency dynamics over time, a process that can be modeled using a variety of approaches. Connecting the expectations from these models with observed genomic data is often achieved using the site frequency spectrum (SFS), a genome-wide summary of genetic polymorphism within and between populations (Sawyer and Hartl 1992; Adams and Hudson 2004; Caicedo *et al*. 2007; Gutenkunst *et al*. 2009; Nielsen *et al*. 2009). The ease and affordability of collecting genomic SNP data make inferences of demography using the SFS especially appealing, highlighting their importance in gaining insights into the historical factors affecting neutral variation in populations. Several recent analyses have also applied SFS-based methods to infer the fitness effects of mutations (Kim *et al*. 2017; Tataru *et al*. 2017; Fortier *et al*. 2019), allowing researchers to model patterns of selection while simultaneously controlling for demography (Williamson *et al*. 2005).

Generating the SFS from a demographic model is a well-studied problem with several possible approaches, all based on different underlying methodologies, currently implemented [e.g., diffusion: Gutenkunst *et al*. (2009); spectral methods: Lukić and Hey (2012); the coalescent: Excoffier *et al*. (2013); and moment closure: Jouganous *et al*. (2017)]. However, these methods generally assume panmixia or random mating within populations, which may not be a realistic assumption for many groups of organisms that are inbred. The reason for this assumption is that the approximations used by these approaches are all built on top of the Wright-Fisher model and rely on the simplicity of its binomial sampling scheme for deriving expectations. The excess of homozygosity caused by inbreeding deviates from binomial expectations, leading to changes in the observed SFS that cannot be captured by models assuming random mating that may affect estimates of demography. Generalizations of the standard Wright-Fisher model have been made to include inbreeding through partial self-fertilization (Wright 1951). Nevertheless, these modifications have yet to be implemented in SFS-based methods for demographic inference.

Despite this lack of available SFS-based methods, previous approaches to infer demography from inbred samples have successfully used alternative representations of genomic data to capture the extent to which samples share blocks of their genome through non-random mating. This typically entails identifying parts of the genome that are identical by descent (IBD), or that contain runs of homozygosity (ROH), and using the length and distribution of these blocks to infer levels of inbreeding and past population size dynamics (Kirin *et al*. 2010; Kardos *et al*. 2017; Browning *et al*. 2018). Large IBD blocks are usually an indication of recent inbreeding, while the frequency and distribution of smaller IBD blocks, which are shared due to common ancestry rather than inbreeding, contain information about more long-term trends in population size (Kirin *et al*. 2010; Ceballos *et al*. 2018). However, these methods are generally only used to model size changes in single populations, which doesn’t allow them to estimate other important demographic events such as population divergence or rates of gene flow. Furthermore, the reliance of these methods on fully sequenced genomes prevents them from being used in systems that lack such resources.

The ability to estimate demography in organisms that do not have a reference genome is a strength of SFS-based methods. This flexibility allows researchers using reduced representation methods (e.g., restriction enzyme-based approaches) to collect genomic data for demographic inference. A large motivating factor for the work that we have conducted here is to understand demography in domesticated crop species, which are often highly inbred due to how they are bred and propagated (Gaut *et al*. 2018). Inbreeding is also of great concern in threatened and endangered species (Shafer *et al*. 2015; Xue *et al*. 2015; Robinson *et al*. 2016, 2019). For many of the most economically or ecologically important species in these categories, full genome sequences are typically available and can be used to guide estimates of genetic variation and past population dynamics that will help to inform breeding practices or management strategies, respectively. However, for less well-studied agricultural or threatened species, it is crucial to have tools available that can also provide this essential information without necessarily needing to obtain a fully sequenced genome.

In this paper we introduce a new method for including inbreeding in estimates of demography by modifying the sampling distribution used to generate the expected SFS for a given demographic model. We have implemented the approach in the Python package ∂*a*∂*i* (Gutenkunst *et al*. 2009), building on top of its existing machinery for estimating demography using the diffusion approximation. To assess our ability to co-estimate inbreeding and demography, we generated frequency spectra in both ∂*a*∂*i* and SLiM (Haller and Messer 2019) and used the new model to make inferences from these simulated data. We also used simulated frequency spectra from ∂*a*∂*i* to see how inbreeding affects estimates of demography when it is ignored. Finally, we used genomic data from two empirical examples, American pumas (*Puma concolor*) and domesticated cabbage (*Brassica oleracea* var. *capitata*), and evaluated estimates of their demographic histories both with and without inbreeding. In general, our model is shown to be accurate even for highly inbred populations (*F* = 0.9). We also found that failing to account for inbreeding leads to inaccurate estimates of parameters and poor model fit. Taken together, the model we have developed provides a powerful tool to jointly estimate inbreeding and demography, and will help to facilitate evolutionary inferences in a wide-range of species.

## New Approaches

We start with a brief overview of the SFS and describe its derivation from the population distribution of allele frequencies (DAF), which can be obtained using the diffusion approximation as described previously (Gutenkunst *et al*. 2009). We then propose a probability model for calculating the number of derived mutations in an inbred population and provide an expression for the expected SFS incorporating this distribution. Using this expression for the expected SFS with inbreeding we can perform parameter inference with a composite likelihood assuming a Poisson Random Field model (Sawyer and Hartl 1992).

### The Site Frequency Spectrum

The site frequency spectrum (SFS) is a multidimensional summary of genetic variation within and across populations that records how often derived biallelic variants of different frequencies are observed in a sample of individuals. For example, given a sample of 20 chromosomes (10 diploid individuals) from three populations, the SFS entry at index [3,8,17] records how often we observe a variant in three, eight, and 17 out of the 20 chromosomes in populations one, two, and three, respectively. In general, for *P* populations with sample sizes *n*_1_, *n*_2_, …, *n*_*P*_, we index the SFS using [*d*_1_, *d*_2_, …, *d*_*P*_] to record how often we observe a variant with frequency *d*_1_, *d*_2_, …, *d*_*P*_ in populations one through *P*.

The observed SFS can be obtained from empirical data by tabulating derived SNP frequencies across sampled populations to generate the *P*-dimensional array described above. When a derived allele cannot be determined, we can instead record the frequency of the minor allele, effectively “folding” the spectrum in half by only considering the variants with frequency less than 0.5. Demographic inference can then be conducted by comparing the observed SFS with the SFS obtained from a demographic model (Sawyer and Hartl 1992).

Given the *P*-dimensional distribution of allele frequencies obtained from a given demographic model, *ϕ*, the expected SFS can be obtained by calculating the probability of drawing *d*_1_, …, *d*_*P*_ derived alleles while integrating across the distribution of allele frequencies in the populations. Within each population, the number of derived alleles has a binomial distribution under panmixia. We then integrate across all possible allele frequencies, weighting the binomial probability of drawing *d*_*i*_ derived alleles by the density determined by *ϕ* within population *i*. Taking this *P*-dimensional integral across the weighted product of binomial probabilities gives us the expression for the joint expected SFS:

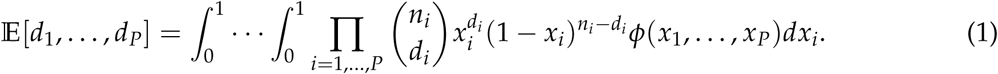

### The Expected SFS with Inbreeding

Through its use of binomial sampling, the preceding derivation for the expected SFS makes the assumption that matings within populations are random. When inbreeding has occurred, individual genotypes are more likely to be homozygous due to being IBD. One way to capture this excess in homozygosity is to incorporate the inbreeding coefficient *F* into a generalized form for the expected genotype frequencies under Hardy Weinberg equilibrium (Wright 1951). Here we use an alternative model that captures the fact that genotypes within populations will be correlated due to inbreeding, pushing the distribution of genotypes towards homozygotes. To capture this correlation among genotypes, Balding and Nichols (1995, 1997) proposed a probability model to incorporate inbreeding using a beta-binomial distribution. Under this model, individual genotypes are a random variable, *G*_*i*_ ∈ {0, 1, 2}, for the number of copies of the derived allele in individual *i* (*i* = 1, …, *n*) such that *Pr*(*G*_*i*_ = *g*) at an individual locus with allele frequency *p* ∈ (0, 1) and population inbreeding coefficient *F* ∈ (0, 1) is beta-binomial with the following form:

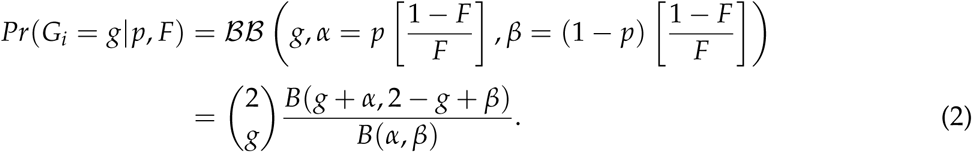

Here ℬ ℬ denotes the probability mass function for the beta-binomial distribution and *B*(*x, y*) is the beta function with dummy parameters *x* and *y*. The parameterization of 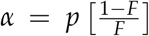 and 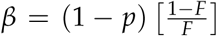 introduces the overdispersion of probability towards homozygous genotypes that is expected as inbreeding increases (Balding and Nichols 1995, 1997).

To get the expected SFS, we need to be able to model the total number of derived alleles sampled in the population, which is the sum across the genotypes of all individuals. Given a sample of *n* diploid individuals (2*n* chromosomes), we use the random variable *D* ∈ { 0, …, 2*n*} to denote this quantity. The probability mass function for *D* is an *n*-fold convolution of beta-binomial distributions, which does not have a simple distributional form. However, we can obtain the probability mass function by considering all possible combinations of the probability of drawing *D* = *d* alleles across *n* beta-binomial distributions, giving us a closed form expression for the convolution of *n* beta-binomial random variables:

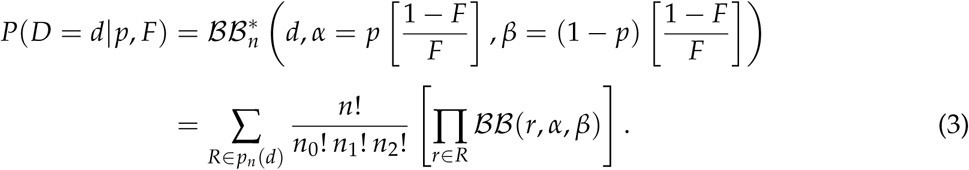

Breaking this down, we can think of it as enumerating all possible ways to generate genotypes in *n* individuals such that they sum to *d*, times the beta-binomial probability of sampling each genotype. More specifically, let *p*_*n*_(*d*) be an array of integer partitions with *n* entries that sum to *d* such that all entries in the partition are 0, 1, or 2 (corresponding to the possible genotype values). For example, the partitions defined by *p*_5_(4) are [2, 2, 0, 0, 0], [2, 1, 1, 0, 0], and [1, 1, 1, 1, 0]. Then for each of these partitions, we use the multinomial coefficient 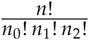, with *n*_0_, *n*_1_, and *n*_2_ corresponding to the number of partition entries equal to 0, 1, and 2, respectively, to account for all possible rearrangements of the partition entries. Next, we multiply the beta-binomial probability for each genotype in a partition using Eq. 2. Taking the product across all possible partitions gives us the full expression for the *n*-fold convolution, which we denote 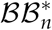 (* is the mathematical operator for convolutions). Inserting this distribution into Eq. 1 gives us the final form for the expected SFS with inbreeding:

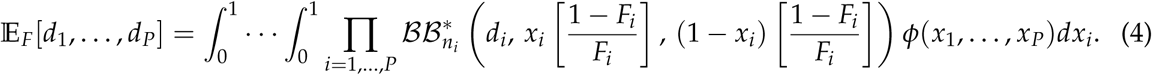

We have written a small R Shiny application illustrating the probability distribution for the beta-binomial convolution (available on GitHub). Figure 1 also shows a sample of example frequency spectra for different levels of inbreeding.

**Figure 1:**
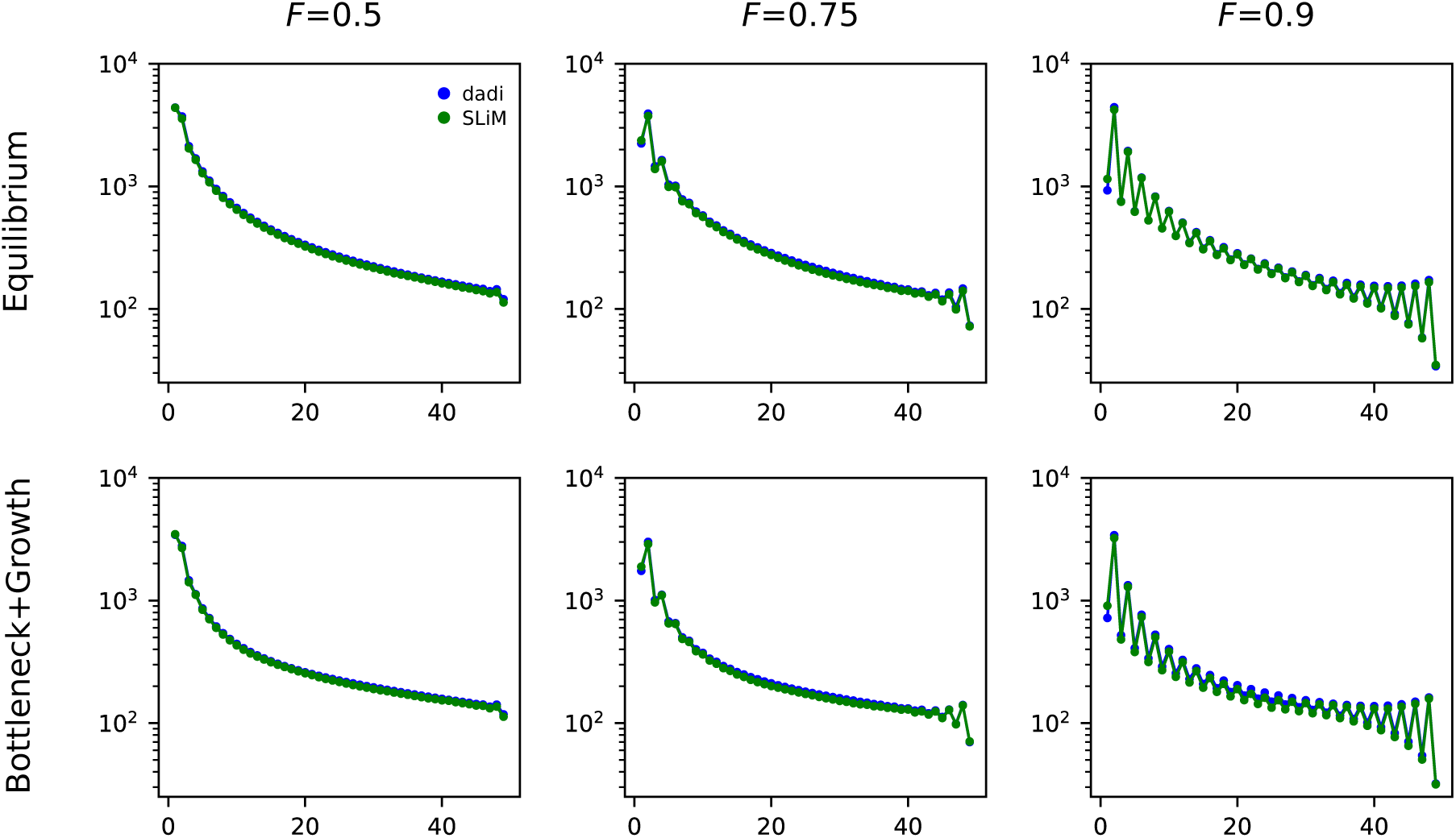
Comparison of expected spectra for *F* = 0.5, 0.75, and 0.9 between ∂*a*∂*i* (blue) and SLiM (green) for the equilibrium and bottleneck+growth models.

## Results

### Comparison with SLiM

We used SLiM (Haller and Messer 2019) to validate the expectations of the SFS with inbreeding by simulating frequency spectra under three models (described in more detail in the **Simulations** section below): a simple equilibrium model (standard neutral model), a one-population bottleneck and growth model, and a two-population divergence and one-way migration model. Inbreeding was assumed to occur through selfing and expected frequency spectra were obtained by taking the mean of 5000 simulations for each model. Figure 1 plots the comparison between the SFS obtained from ∂*a*∂*i* (blue) and SLiM (green) for the equilibrium and bottleneck models with *F*=0.5, 0.75, and 0.9, respectively. The frequency spectra for these models for *F*=0.1 and *F*=0.25 are presented in Figure S1 and the comparisons for the two-population divergence model are in Figure S2. The percent differences between the frequency spectra from ∂*a*∂*i* and SLiM were between 0.1% to 0.2% for the one-population models and were between 0.02% to 0.03% for the two-population model, demonstrating that our results from modeling the expected SFS with beta-binomial distributions corresponds well with the spectra simulated from SLiM.

We also used simulated frequency spectra from SLiM to estimate parameters for these three models in ∂*a*∂*i*. Figure 2 shows the distribution of estimated inbreeding coefficients for the bottleneck and growth model (RMSD = 0.094) and divergence and one-way migration models (RMSD = 0.163). Similar plots for all other estimated parameters across all three models are presented in Figures S3–S5.

**Figure 2:**
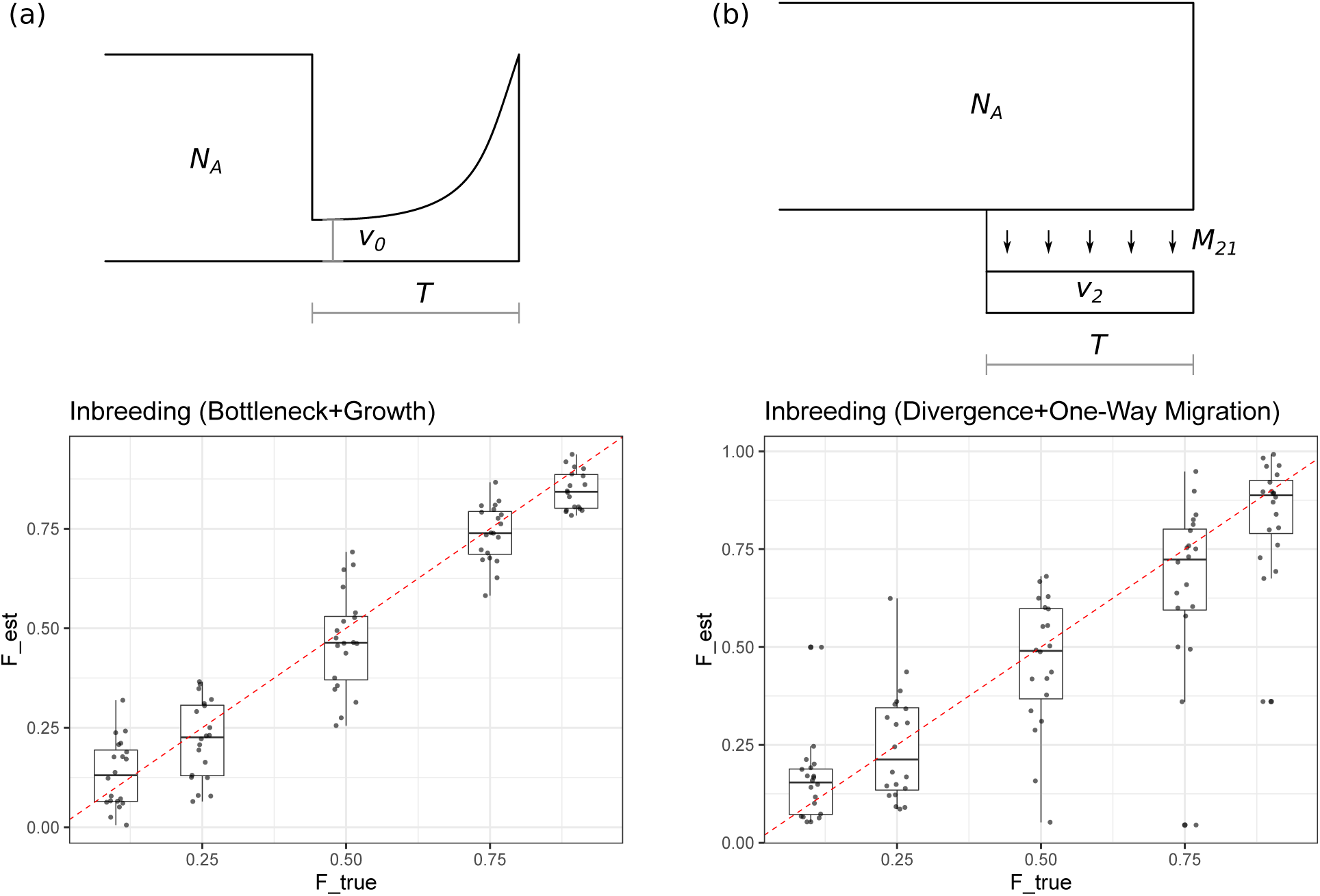
(a) Estimates of *F* from data generated with SLiM for the bottleneck and growth model (lower) plus an illustration of the model (upper). In this model, *N*_*A*_ is the ancestral population size, *ν*_0_ is the size of the bottleneck (proportion of *N*_*A*_ remaining after population reduction), and *T* is the amount of time for the population to recover back to a size of *N*_*A*_. (b) Estimates of *F* from data generated with SLiM for the divergence with one-way migration model (lower) plus an illustration of the model (upper). *N*_*A*_ in this model is the same as the bottleneck model, *ν*_2_ is the size of the diverging population (again a proportion of *N*_*A*_), *T* is the divergence time between populations, and *M*_21_ is the one-way migration rate of individuals from population one into population two.

### Simulations

#### Simulation 1: Co-Estimating Inbreeding and Demography

To assess our ability to estimate demographic parameters under increasing levels of inbreeding (*F*= 0.1, 0.25, 0.5, 0.75, and 0.9), as well as the inbreeding coefficient within a population itself, we performed demographic inference using simulated frequency spectra under three models: (1) a standard neutral model, (2) a one-population bottleneck and growth model, and (3) a two-population divergence model with unidirectional gene flow (models two and three are illustrated in Figure 2). For the standard neutral model, the inbreeding coefficient is the only parameter that needs to be estimated. The one-population bottleneck and growth model has three parameters: the inbreeding coefficient, the relative size of the bottlenecked population (*ν*_0_ = 0.1, 0.25, and 0.5), and the recovery time back to the original size (*T* = 0.1, 0.2, and 0.3). The two-population model has four parameters: the inbreeding coefficient, the relative size of the diverging population (*ν*_2_ = 0.1, 0.25, 0.5), the time of divergence from the main population (*T* = 0.1, 0.2, and 0.3), and the rate of gene flow from the main population into the diverged population (*M*_21_ = 0.5, 1.0, and 1.5). All parameters are specified relative to the ancestral population size, which in ∂*a*∂*i* defaults to 1.0.

Figures S6–S8 shows the distribution of estimated inbreeding coefficients across 20 replicate experiments for every combination of simulation parameters for the equilibrium, bottleneck, and divergence models. For all three models, we are able to recover accurate estimates of *F* (Model 1 RMSD: 0.0139; Model 2 RMSD: 0.0176; Model 3 RMSD: 0.0406) even when inbreeding is quite high (*F* = 0.9). Figure S7 also shows plots for estimates of bottleneck size and recovery time across inbreeding levels for model two. The RMSD for these estimates across all simulated values were 0.0236 and 0.0184 for *ν*_0_ and *T*, respectively. Figure S8 shows similar plots for estimates of population size, divergence time, and one-way migration rate across inbreeding levels for model three. The RMSD for these estimates across all simulated values were 0.0131 for *ν*_2_, 0.0103 for *T*, and 0.158 for *M*_21_.

#### Simulation 2: Parameter Estimation When Inbreeding is Ignored

To understand the impact of ignoring inbreeding on demographic inference, we simulated data sets with inbreeding under the same bottleneck and divergence models as above (models two and three) but performed inference under the assumption that inbreeding was absent. Because of initial issues with convergence in these analyses, particularly with the bottleneck model, and the fact that higher levels of inbreeding cause increasingly conspicuous changes to the SFS (e.g., see Figure 1), we used a smaller range for *F* in these simulations: 0.1, 0,2, 0.3, 0.4, and 0.5.

Parameter estimates for the bottleneck model had higher rates of error compared to when inbreeding was directly modeled. The RMSDs for *ν*_0_ and *T* were 0.191 and 0.117, respectively. Estimates of these parameters also got worse as inbreeding increased (Figure S9), clearly demonstrating the issues that can arise when inbreeding is ignored. In contrast, results for the divergence model were surprising in that they didn’t show the high levels of estimation error seen with the bottleneck model (Figure S10). The RMSD values for the parameters of the divergence model were 0.0261 for *ν*_2_, 0.0130 for *T*, and 0.142 for *M*_21_. Interestingly, the RMSD for *M*_21_ was actually lower in this simulation experiment than when inbreeding was modeled (0.158). However, the increase in RMSD for the simulations where inbreeding is modeled is due to using higher levels of inbreeding (*F* > 0.5). If we restrict the calculation of RMSD in the estimates including inbreeding to only those with *F* ≤ 0.5, the RMSD is lower than when inbreeding is ignored, as expected (0.109). RMSD values for *ν*_2_ and *T* where higher for model two than in *Simulation 1*, indicating that these parameters may be more sensitive to the effects of unmodeled inbreeding.

#### Simulation 3: Masking Rare Variants

Several techniques to ‘side-step’ the impact of inbreeding have been taken in empirical analyses. This includes sampling only a single chromosome per site, per individual (e.g., Beissinger *et al*. 2016; Koenig *et al*. 2019) or masking rare variants (e.g., Cornejo *et al*. 2018), which are disproportionately affected at lower levels of inbreeding (Figure 1). Since sampling only a single chromosome cuts the sample size in half, and investigations on the effect of sample size on demographic inference have already been explored (Robinson *et al*. 2014), we instead focused on the effect of masking rare variants under increasing levels of inbreeding. For the bottleneck model we masked the singleton and doubleton entries of the 1D-SFS, and for the divergence model we masked the bottom corner of the 2D-SFS (ie singletons, doubletons, and their combinations across both populations). We then used the same range of parameters as in the previous simulations to see how much masking affected our inferences.

For the bottleneck and growth model, data masking had a small but noticeable effect on parameter estimation. The bottleneck size was estimated with less accuracy compared to when inbreeding was included (RMSD = 0.0296) and estimates of recovery time also had higher error (RMSD = 0.0218), typically in the direction of underestimation (Figure S11). The effect of masking was more pronounced in the divergence model (Figure S12), particularly for the migration parameter, where the amount of gene flow was almost always underestimated across all parameter combinations (RMSD = 0.193). Estimates of population size and divergence time were also slightly underestimated when compared to models including inbreeding (RMSD = 0.0122 and 0.0103, respectively) but the effect was less pronounced.

#### Simulation 4: Misspecified Inbreeding

As a final test of the model for inbreeding, we simulated frequency spectra under the bottleneck and divergence models without inbreeding but included it as a parameter to be estimated. The expectation in this case is that inbreeding should be estimated close to 0 and that its inclusion in the model does not lead to poor estimates of other model parameters. However, for both models, the inbreeding parameter was always estimated to be greater than 0. The mean estimates of *F* for the bottleneck and divergence models were 0.0934 and 0.212, respectively. Nevertheless, despite not estimating an absence of inbreeding, the other model parameters were estimated with only slightly higher levels of error (Figures S13 and S14). For the bottleneck model, bottleneck size and duration had RMSD values of 0.0280 and 0.0268, respectively, which are both higher levels of error than the simulations where inbreeding was truly present. Parameters in the divergence model had RMSDs of 0.0183 for *ν*_2_, 0.0110 for *T*, and 0.132 for *M*_21_, showing that the two-population model was not strongly affected by the level of inbreeding estimated in population two.

### Empirical Examples

#### American Puma

The American puma (*Puma concolor*) is an iconic carnivore distributed primarily in western North America and South America, occupying a large diversity of habitats across its range. However, in the eastern United States, the only remnant population is the highly endangered Florida panther (Hansen 1992; Culver *et al*. 2000). Florida panthers have been the subject of large-scale conservation efforts aimed at ameliorating the adverse effects of small population size, including moving individuals from their closest sister population, the Texas puma, to introduce novel genetic variation (Seal and Lacy 1994; Johnson *et al*. 2010). Using genomic data from five individuals of Texas pumas and two individuals of ‘canonical’ Florida panthers from Ochoa *et al*. (2019), we estimated the demographic history of these two populations to investigate their divergence time, changes in population size, and levels of inbreeding (see cartoon in Figure 3). More specifically, we fit a model that included an initial change in population size to mimic the colonization of North America by the Texas population (*N*_*TX*_), the duration of time spent at the new population size (*T*_1_), the divergence time between Texas pumas and Florida panthers (*T*_2_), and the inbreeding coefficients for both the Texas and Florida populations (*F*_*TX*_ and *F*_*FL*_).

**Figure 3:**
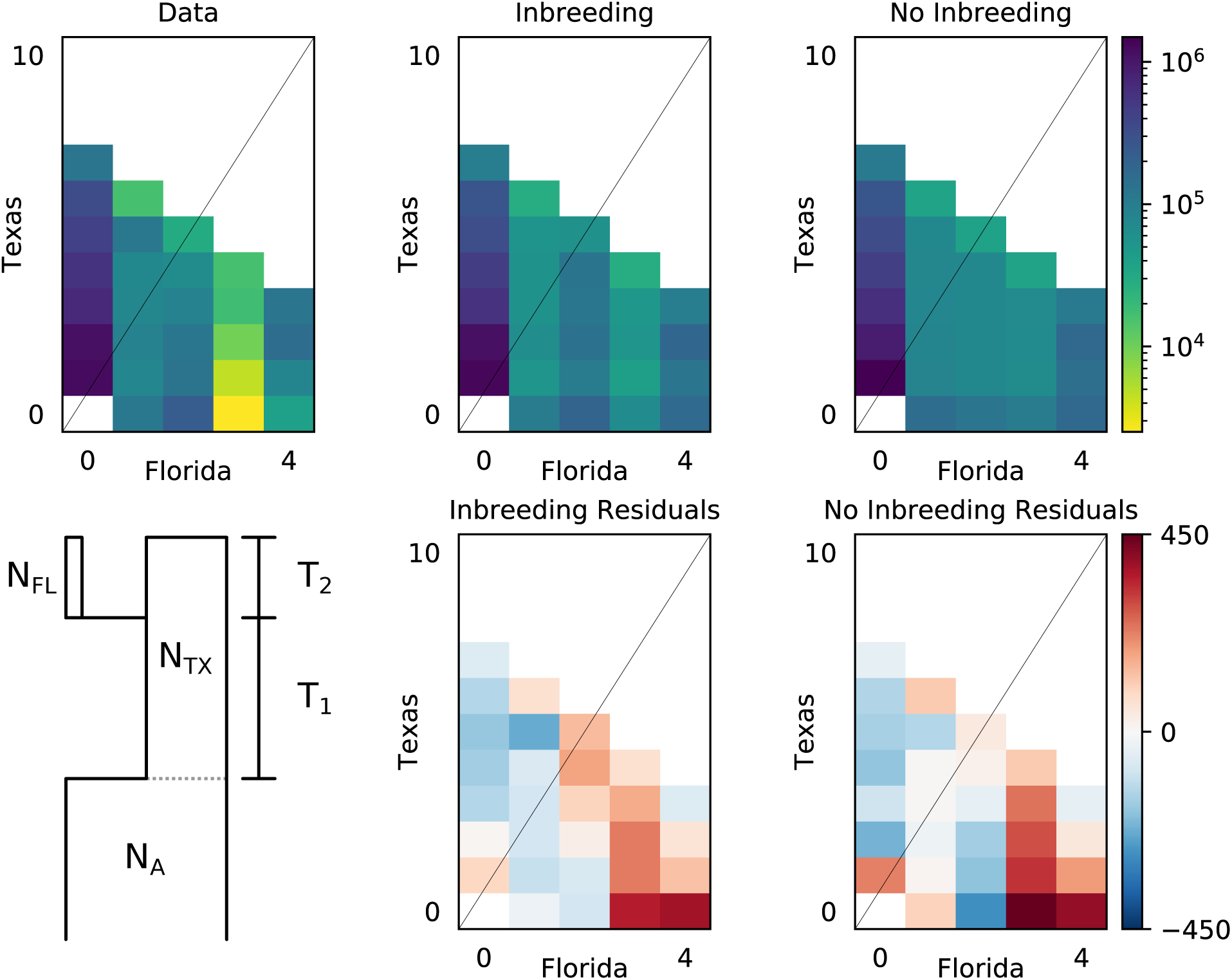
The observed joint site frequency spectrum for *Puma concolor* in Texas and Florida, along with the model fit and residuals, for models with inbreeding (middle) and without inbreeding (right). Residuals for each model are plotted below their expected spectra and a cartoon representation of the proposed demographic model is given in the bottom left.

After processing (see **Methods**), 6,262,417 variant sites were retained for constructing the 2D-SFS. Because we lacked a suitable outgroup for determining ancestral versus derived allelic states, we used the folded SFS for all model fits. Table 1 lists parameter estimates and their 95% confidence intervals for models fit with and without inbreeding (*ϵ* = 10^−2^) and uncertainty estimates across different step sizes for numerical differentiation using the Godambe Information Matrix (Coffman *et al*. 2015) are presented in Tables S3 and S4. In both models, the Texas and Florida populations are estimated to have diverged 7,000–8,000 years ago, with both also having similar estimates of the ancestral population size (120,000–130,000 individuals). As expected, the Florida population experienced a severe reduction in population size down to 1,200–1,600 individuals, as well as having a high estimate of *F* in the model including inbreeding (*F*_*FL*_ = 0.607). Texas pumas were also inferred to be inbred, though less so than Florida panthers (*F*_*TX*_ = 0.440). Estimates of population size for the Texas population were different between the models with and without inbreeding (70,800 individuals versus 23,700 individuals) and the duration of the initial population size change (*T*_1_) were especially different as well (247,000 years versus 26,800 years). The log-likelihoods for the model with and without inbreeding are − 318058.079 and − 453003.048, respectively, and the Godambe-adjusted likelihood ratio statistic is 425.489 (p-value = ∼ 0.0; Coffman *et al*. 2015), demonstrating that the model with inbreeding has a significantly better fit to the data. In addition, when comparing the predicted SFS from the models with the observed SFS (Figure 3), the residuals for the model with inbreeding were lower overall, providing even more support for preferring the model with inbreeding. Uncertainty estimates were also typically more stable across step sizes for the model with inbreeding.

**Table 1:**
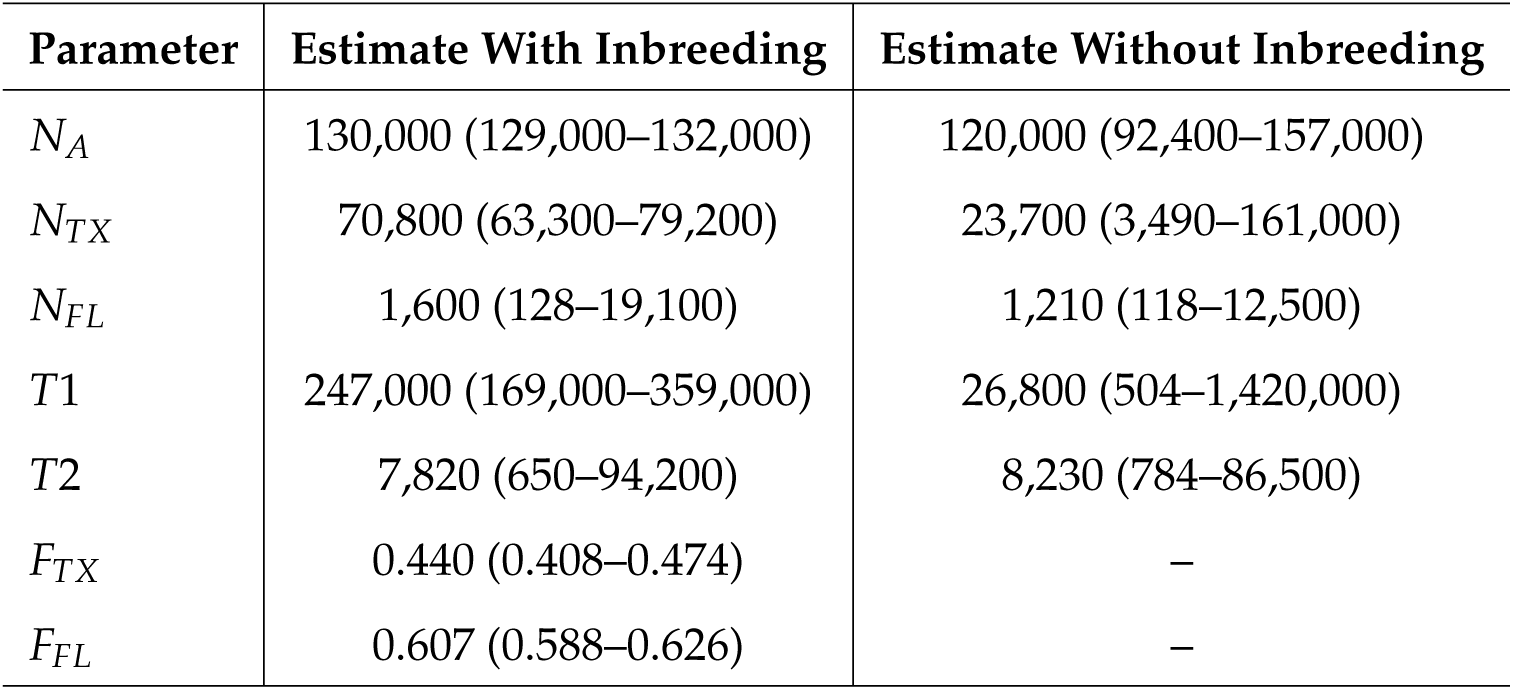
Parameter estimates for *Puma concolor* from demographic models estimated both with and without inbreeding. 95% confidence intervals are given in parentheses and were estimated using a step size of *ϵ* = 10^−2^ for numerical differentiation. Population sizes are given in number of individuals and divergence time is given in years.

#### Domesticated Cabbage

*Brassica oleracea* is an agronomically important plant species cultivated primarily in Europe, Asia, and North America (Maggioni 2015). It is especially well-known for its morphological diversity, having been domesticated into several different crops including broccoli, Brussels sprouts, cauliflower, cabbage, kale, and kohlrabi, among others. The timing and origin of domestication for these different *B. oleracea* crops is still uncertain, but several hypotheses have been proposed to explain their evolutionary history (Maggioni 2015). Cabbage, *B. oleracea* var. *capitata*, is thought to have been domesticated roughly 500 years ago in the Mediterranean region (Cheng *et al*. 2016a,b), providing an interesting hypothesis that we can test using demographic models.

To infer the demographic history of domesticated cabbage, we used SNP data from publicly available resequencing data for 45 individuals from Cheng *et al*. (2016a,b). We then fit a demographic model for cabbage domestication that included two changes in population size (*N*_1_ and *N*_2_), the amount of time spent at these population sizes (*T*_1_ and *T*_2_), and the level of inbreeding (*F*) [see cartoon in Figure 4]. We used 2,941,018 intergenic SNPs to build the folded SFS for *B. oleracea* var. *capitata* and fit models with and without inbreeding. Parameter estimates were obtained using newly implemented optimization routines in the ∂*a*∂*i* library built on top of the nlopt Python package (Johnson 2014). Parameter estimates and their 95% confidence intervals are listed in Table 2 (*ϵ* = 10^−2^). Uncertainty estimates across different step sizes for numerical differentiation using the Godambe Information Matrix (Coffman *et al*. 2015) are presented in Tables S3 and S4.

**Table 2:** Parameter estimates for *B. oleracea* var. *capitata* from demographic models estimated both with and without inbreeding. 95% confidence intervals are given in parentheses and were estimated using a step size of *ϵ* = 10^−2^ for numerical differentiation. Population sizes are given in number of individuals and times are given in years. Parameters estimated at the upper/lower bound of the given search space are marked with an asterisk (*).

| Parameter | Estimate With Inbreeding | Estimate Without Inbreeding |
| --- | --- | --- |
| $N_A$ | 17,500 (16,900–18,100) | 19,100 (18,500–19,800) |
| $N_1$ | 31,600 (28,900–34,700) | 123,000 (80,400–190,000) |
| $N_2$ | 215,000 (4,910–9,370,000) | 592 (547–641) |
| $T_1$ | 16,600 (12,900–21,200) | 5,870 (5,200–6,620) |
| $T_2$ | 322 (94.2–1,097) | 38.3 (32.5–45.1)* |
| $F$ | 0.578 (0.557–0.599) | – |

**Figure 4:**
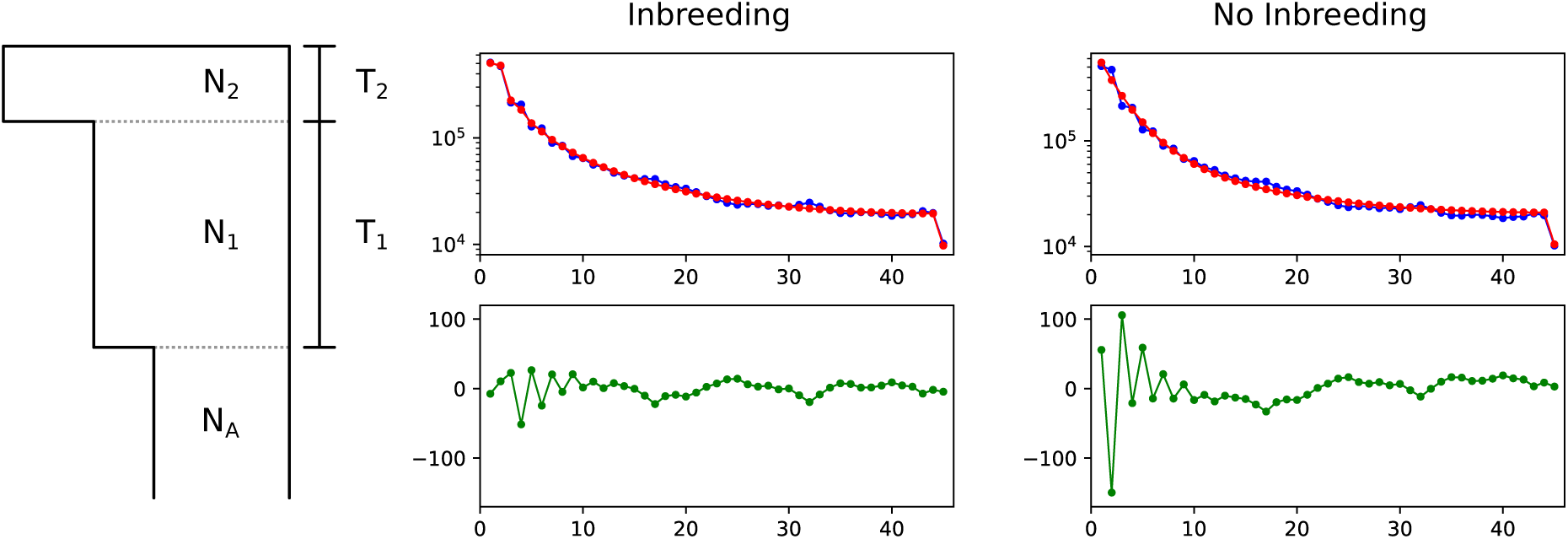
The observed site frequency spectrum for *Brassica oleracea* var. *capitata*, along with the model fit (red) and residuals (bottom panels), for models with inbreeding (middle) and without inbreeding (right). On the left is a cartoon of the proposed demographic model with parameters labeled.

Much like the models inferred with and without inbreeding for American pumas, the estimates of demography for cabbage are markedly different between the two analyses. When inbreeding was not modeled, we infer an ancestral population size for cabbage of 19,100 individuals, which expanded to a size of 123,000 individuals ∼ 6,000 years ago. This expanded population then experienced a very recent and severe bottleneck 38 years ago down to a size of 592 individuals. The time estimate for the bottleneck consistently hit the lower bound of the parameter search space, however, suggesting that this estimate is likely not very reliable. Parameter estimates for the inbreeding model inferred an ancestral population size of 17,500 individuals, which expanded to a size of 31,600 individuals ∼ 17,000 years ago. This population then experienced an even larger expansion to a size of 215,000 individuals 322 years ago. The model with inbreeding inferred *F* to be 0.578, showing that inbreeding in these cabbage samples is fairly high. The log-likelihoods for the model with and without inbreeding were − 4281.145 and − 24330.403, respectively, and the Godambe-adjusted likelihood ratio statistic was 127.562 (p-value = ∼ 0.0; Coffman *et al*. 2015). Figure 4 also shows the observed and predicted SFS for each model plus their residuals. The residual plots clearly show that the model with inbreeding is able to capture more of the ‘zig-zagging’ pattern of the lower frequency variants than the model without inbreeding, demonstrating its overall better fit. Uncertainty estimates were again typically more stable across step sizes for the model with inbreeding.

## Discussion

The prevalence of inbreeding in nature, especially among plant lineages and small and endangered populations, make it an important process to include in demographic models. Unlike previous approaches that rely on full genome sequences to characterize patterns of identity by descent or the distribution of runs of homozygosity, our model uses the frequency spectrum of biallelic SNPs to infer demography, allowing it to be employed not only in model systems but in organisms that lack a suitable reference genome as well. The impact of inbreeding on the SFS has important consequences for demographic inference, however, a result that is well-demonstrated by our simulations and example analyses. The relationship between inbreeding and population size is especially relevant for understanding inferences of past population dynamics. Below we describe this connection in the context of our simulations and the results of our empirical analyses, drawing on previous theoretical work to help qualify our results. We then discuss the importance of considering how our current model behaves for recent versus sustained inbreeding.

### The Effects of Inbreeding on Estimates of Demography

#### Comparison with SLiM

In our analysis of frequency spectra from SLiM, we found a high level of agreement between the expected SFS from the diffusion approximation and beta-binomial model in ∂*a*∂*i* and the mean SFS from the three models we tested in SLiM. In addition, we were generally able to get accurate estimates for the parameters of the three models, though there was a large amount of variation. Part of this is likely driven by only simulating a 1 Mbp region, which limits the number of SNPs being used to build the SFS. A more important contributor to the variation in parameter estimates is the impact of inbreeding itself on the scaling of population level parameters such as *θ*. Previous work in both the diffusion (Pollack 1987) and coalescent (Nordborg and Donnelly 1997; Nordborg 2000) frameworks have derived the appropriate scaling of population-level parameters for inbred populations. In both cases, the equilibrium *θ* of a randomly mating population simply needs to be rescaled by 1 + *F* to obtain the corresponding process with inbreeding (here inbreeding is achieved through selfing). The same rescaling applies to parameters estimated by ∂*a*∂*i* when inbreeding is included, so the appropriate scaling can be achieved by rescaling the affected parameters by 1 + *F* using the estimated value of the inbreeding coefficient.

#### Simulations with ∂a∂i

From our more detailed simulation experiments, we were able to characterize several scenarios where inbreeding adversely affects inferences of demography. For the single-population bottleneck model in particular, not accounting for inbreeding had a dramatic impact on the accuracy of estimated population size. The primary reason for this is that inbreeding, much like population growth or contraction, affects the low frequency entries of the SFS in such a way that these factors are likely confounded (Figure 1).

The result of ignoring inbreeding for the two-population divergence with one-way migration model was much less drastic (Figures S10), such that parameter estimates were often nearly as accurate as when we included inbreeding in the model. It should be noted that in this case the highest level of inbreeding was *F* = 0.5, compared to the highest level in the co-estimating inbreeding simulations (*F* = 0.9). However, the fact that the results did not show the same pattern of extremely poor parameter inference as the one-population model despite also having a bottleneck was noteworthy. One possible explanation for this is that jointly modeling the demography of the inbred, bottlenecked population with the main, non-inbred population provided more information in the 2D-SFS to estimate parameters. Nevertheless, despite having more overall accuracy than the one-population model, parameter estimates in the two-population model were increasingly underestimated with higher levels of inbreeding, demonstrating its adverse effects even when including a second panmictic species in the model.

The other two simulation experiments, masking rare variants and misspecifying inbreeding, provide further examples of the extent to which the variants with lowest frequency are confounded with inbreeding in ways that affect demographic inference. Masking the singleton and doubleton entries of the 1D- and 2D-SFS for the bottleneck and divergence models, respectively, had only a small effect on estimates of population size and the timing of demographic events, showing that the signal for these inferences is also contained in the remaining entries of the SFS. However, estimates of gene flow were consistently underestimated in the divergence model, likely due to the fact that the influx of migrant alleles at low frequency were being masked. Simulations that modeled inbreeding when it was absent provide a different view on the inference of inbreeding and demography. In this case, the inbreeding coefficient was inferred to be ∼ 0.1 and ∼ 0.2 in the one- and two-population models, respectively, even though there was no inbreeding (Figure S13 and S14). The accuracy of the remaining parameters was fairly high; however, there were instances of certain parameter combinations leading to over- and underestimation of the true parameter value. Therefore, to prevent poor estimation of other parameter values, it is advised that inbreeding be included in a model only when there is an observable excess in homozygosity.

#### Results from Empirical Analyses

The impact of inbreeding on the results of our empirical analyses demonstrate the importance of directly estimating this parameter when inferring demography. Analyses with and without inbreeding provided different estimates of population size and duration for Texas pumas and infer strikingly different population size changes during the history of domesticated cabbage. In the case of the American puma, our estimates of population divergence time between Texas and Florida, the timing of movement from South America into North America, reductions in population size (especially for the Florida population), and a high level of inbreeding in Florida panthers are all consistent with previous work (Culver *et al*. 2000; Ochoa *et al*. 2017, 2019).

The demographic history inferred for cabbage provides yet another example of how inbreeding and population contraction can be confounded since estimates of current population size without inbreeding were ∼ 600 individuals, an unrealistic estimate given the prevalence of cabbage cultivation, as well as the clear discrepancy between model fit and the observed SFS for low frequency variants (Figure 4). Including inbreeding, however, provides a potentially revealing look into the domestication history of cabbage, especially regarding the signal for the textbook “domestication bottleneck” (Gaut *et al*. 2018). The expansion of the ancestral cabbage population ∼ 17,000 years ago coincides with the end of glaciation in Europe and, in particular, the Mediterranean region (Hughes *et al*. 2006; Clark *et al*. 2009; Hughes and Woodward 2017). Previous work has also placed the timing of domestication for the cabbage morphotype of *B. oleracea* at approximately 500 years ago (Cheng *et al*. 2016b), which roughly agrees with the date that we estimated for the secondary population expansion. This series of population expansions differs quite conspicuously when compared to what is often expected for domesticated species (e.g., severe bottlenecks; Doe-bley *et al*. 2006; Meyer and Purugganan 2013; Gerbault *et al*. 2014; Gaut *et al*. 2018). Given the relatively high inbreeding coefficient estimated for cabbage (*F* = 0.58), and the fact that ignoring inbreeding led to inferences of a very recent and severe bottleneck, it is possible that past inferences of domestication bottlenecks have been partially misled by the occurrence of inbreeding when inferring population dynamics.

### Short-Versus Long-Term Inbreeding

In a review on the effects of inbreeding, Deborah Charlesworth (2003) discussed the temporal aspects of its impact on genetic diversity, distinguishing between the short-term consequences on patterns of diversity (i.e., excess homozygosity compared to panmixia) versus the long-term effects of inbreeding that lead to an overall reduction in the effective size of the population. The method we have introduced here is capable of modeling inbreeding in both categories by not only fitting the physical manifestation of inbreeding in the SFS (i.e., spikiness), but also by being able to appropriately scale the diffusion process to account for the reduction in diversity caused by inbreeding 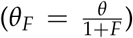. The reduction in effective population size, as well as the recombination rate, that inbreeding causes has important consequences for the impact of selection and the rate of adaptation in inbred populations (Charlesworth 1992; Hartfield and Glémin 2016; Hart-field and Bataillon 2019). Therefore, studies aiming to identify the targets of selection in inbred, non-equilibrium populations must exercise special caution. This is especially relevant for domesticated species and organisms of conservation concern, whose evolutionary histories can often involve drastic changes in population size. Moving forward, the joint inference of demography, inbreeding, and selection will be an important advance for better understanding their collective contributions to genetic variation, as well as having potentially large consequences on informing decision making in agriculture and the designation of protection status for threatened or endangered species.

## Materials and Methods

### Comparison with SLiM

Simulations to validate the expected SFS with inbreeding were conducted using SLiM v3.3 (Haller and Messer 2019). To include inbreeding, we set the selfing rate in SLiM to *s* = 2*F*/(1 + *F*), *F* = 0.1, 0.25, 0.5, 0.75, and 0.9, and conducted 50 independent simulations, randomly sampling 25 individuals with replacement 100 times for each replicate, for a total of 5000 simulated spectra for each of three models: (1) equilibrium/standard neutral model, (2) bottleneck and growth model, and (3) divergence with one-way migration model (models two and three are depicted in Figure 2). Each simulation used *θ* = 4*N*_*A*_ *Lµ* = 10, 000, with *N*_*A*_ = 1000, *L* = 1 × 10^6^ bp, and *µ* = 2.5 × 10^−5^. We also set the recombination rate, *r*, equal to the mutation rate. For all models, we simulated 10,000 generations of burn-in to allow the ancestral population to reach equilibrium and included selfing from the start of the simulation for models one and two. Individuals were sampled directly after this phase for the standard neutral model. For model two, after burn-in, we reduced the population size to 250 individuals and then allowed the population size to recover exponentially back to 1000 individuals over 400 generations: 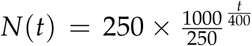. Model three started with an outcrossing equilibrium population, from which we split off a selfing population with a size of 250 individuals. These two populations were then simulated forward for an additional 400 generations with the selfing population receiving migrants from the original population at a rate of *m*_21_ = 5 × 10^−4^. At the end of each simulation, individual genotype information was exported in variant call format and summarized using a Python script to obtained the SFS (available on GitHub). The expected SFS was then calculated by taking the mean value of each entry of the simulated spectra across replicates for each model and each value of the inbreeding coefficient. This simulation routine was also replicated to generate 20 independent data sets for each model across the five levels of inbreeding to infer parameters using ∂*a*∂*i* v2.0.3 (Gutenkunst *et al*. 2009). Models were specified in Python v3.7 using the parameterizations described above and depicted in Figure 2. Parameters were estimated for each simulated frequency spectrum using 100 optimization runs initiated from different random starting points. Parameter estimates with the highest log-likelihood were then recorded for comparison with the true simulated values using the root mean squared deviation (RMSD).

Because inbreeding rescales the effective population size by a factor of 1 + *F* (Pollack 1987; *Nordborg and Donnelly 1997), and* ∂*a*∂*i* estimates parameters relative to the ancestral population size, we rescaled parameters in ∂*a*∂*i* in the following ways for the simulations above. For the standard neutral model, we included selfing from the start of the simulation, so for our comparison between SLiM and ∂*a*∂*i* we divided the original *θ* of 10,000 by 1 + *F*. For the bottleneck and growth models, the ancestral population was also inbred, so we rescaled *θ* by again dividing by 1 + *F*. The recovery time for this model was always set to 400 generations in SLiM, which in ∂*a*∂*i*’s units would be equal to 0.2 × 2*N*_*A*_. Because the effective ancestral population size gets smaller as inbreeding increases, we had to account for this by multiplying by a factor of 1 + *F*. However, when inferring parameters under this model, we have to rescale in the opposite direction by dividing by 1 + *F* to get the correct estimate for the number of generations relative to the ancestral population size. Finally, for the divergence with one-way migration model, the ancestral population is not inbred, so the only rescaling that needs to be done is for the size of the diverged selfing population (0.25*N*_*A*_ in ∂*a*∂*i* units). When comparing the expected SFS between ∂*a*∂*i* and SLiM, we divide 0.25 by 1 + *F* to get the correct size for the inbred population. When inferring parameters, we instead multiply by 1 + *F* to recover the original 0.25*N*_*A*_. In practice, it should be possible to estimate models assuming an outcrossing ancestral population and including a change in population size to account for the effects of inbreeding.

### Simulations

Simulations to explore a greater breadth of parameters were conducted in Python 3.7 using functions available in the ∂*a*∂*i* library (v2.0.3). For each simulation experiment, we used the same basic setup for simulating frequency spectra under the two main models that were tested. The two models were (1) a single-population model experiencing a bottleneck of varying size [0.1*N*_*A*_, 0.25*N*_*A*_, 0.5*N*_*A*_] followed by exponential growth over different time scales [0.2*N*_*A*_, 0.4*N*_*A*_, 0.6*N*_*A*_] back to the original size and (2) a two-population model where a small subpopulation diverges from the main population at different times in the past [0.2*N*_*A*_, 0.4*N*_*A*_, 0.6*N*_*A*_], going through a bottleneck of different sizes [0.1*N*_*A*_, 0.25*N*_*A*_, 0.5*N*_*A*_] and receiving migrants from the main population at different rates [0.25/*N*_*A*_, 0.5/*N*_*A*_, 0.75/*N*_*A*_]. For the **Co-Estimating Inbreeding and Demography** simulations, we generated SFS under a standard neutral model with inbreeding as well. We also used a larger range of inbreeding coefficients for this experiment (*F*_*IS*_=0.1, 0.25, 0.5, 0.75, and 0.9). The remaining three simulations that were not focused on estimating inbreeding used a smaller range (*F*_*IS*_=0.1, 0.2, 0.3, 0.4, and 0.5) since optimizations at higher inbreeding levels generally failed to converge. Each simulation experiment was replicated 20 times, with each replicate having 25 individuals sampled per population and running 50 independent optimizations. Site frequency spectra were generated for each replicate by first getting the expected SFS for the model with the true parameters, followed by scaling the SFS using *θ* = 10, 000 and sampling chromosomes assuming a Poisson distribution (`sample()` method in the `Spectrum class` within ∂*a*∂*i*). Parameter estimates with the highest log-likelihood were selected from the 50 optimization runs for each replicate.

We evaluated parameter estimates for each experiment (including estimates with SLiM above) by comparing the estimated values with the true parameters by calculating the RMSD in R v3.6.1 using the tidyverse package v1.2.1 (R Core Team 2019; Wickham *et al*. 2019). Plots from R were generated using ggplot2 v3.2.1 (Wickham 2009). Plots from Python were made using matplotlib v3.1.1 (Hunter 2007) or plotting functions within the ∂*a*∂*i* library.

### Empirical Examples

#### American Puma

Genome-wide variant data from Ochoa *et al*. (2019) were obtained from the authors for five Texas pumas and two Florida panthers in variant call format (VCF). SNPs within annotated genes were removed using bedtools v2.28.0 (Quinlan and Hall 2010), followed by processing with VCFtools v0.1.16 (Danecek *et al*. 2011) to retain only biallelic SNPs with no missing data. The final data set contained 6,262,417 sites, which we converted from VCF format into ∂*a*∂*i*’s ‘SNP data format’ using a Python script (available on GitHub) for demographic analysis. We then estimated demographic parameters in ∂*a*∂*i* using 100 independent optimization runs from different random starting points (Gutenkunst *et al*. 2009). Parameters were converted from estimated ratios of the ancestral population size (*N*_*A*_) to real units using a mutation rate of *µ* = 2.2 × 10^−9^, a generation time of 3 years, and a sequence length of 2,564,692,624 bp Ochoa *et al*. (2019). Confidence intervals were estimated using the Godambe Information Matrix with 100 bootstrapped frequency spectra that were constructed by randomly sampling genome scaffolds with replacement until we reached the same number of scaffolds as the original full genome (Coffman *et al*. 2015). The Godambe Information Matrix uses numerical differentiation to estimate uncertainty and requires a step size (*ϵ*) to be chosen. The choice of step size should be roughly proportional to the size of the uncertainty that is being estimated. To evaluate which step size was most appropriate for American pumas, we used the bootstrapped spectra to estimate uncertainties across a range of step sizes: [10^−2^ – 10^−7^ by factors of 10]. These bootstrapped frequency spectra were also used to conduct a likelihood ratio test between the models with and without inbreeding using the `LRT adjust()` method in ∂*a*∂*i* and comparing the test statistic to a weighted sum of *χ*^2^ distributions with zero, one, and two degrees of freedom (Ota *et al*. 2000):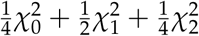. This weighted sum is used because we are testing whether the inbreeding coefficients for the Texas and Florida populations are equal to 0, which is the lower boundary of their parameter space since we are assuming inbreeding coefficients cannot be negative. Because of this, the typical normality assumptions used in the construction of the likelihood ratio test do not apply and we must adjust the distribution being used for assessing the significance of the likelihood ratio test statistic (Ota *et al*. 2000).

#### Domesticated Cabbage

We obtained a VCF file containing SNP calls for 45 cabbage individuals from resequencing data in Cheng *et al*. (2016a,b). We then filtered out genic SNPs with bedtools v2.28.0 using gene annotations from http://www.genoscope.cns.fr/externe/plants/chromosomes.html (Belser *et al*. 2018). Biallelic SNPs containing no missing data were extracted with VCFtools v0.1.16 for a final data set with 2,941,018 variable sites. Demographic parameters were estimated in ∂*a*∂*i* with the BOBYQA algorithm implemented in the nlopt Python package using 100 independent optimization runs from random starting points (Gutenkunst *et al*. 2009; Powell 2009; Johnson 2014). Parameters were converted from estimated ratios of the ancestral population size to real units using a mutation rate of *µ* = 1.5 × 10^−8^, a generation time of 1 year, and a sequence length of 411,560,319 bp (chromosomes minus genic regions). Confidence intervals were then estimated using the Godambe Information Matrix with 100 bootstrapped frequency spectra that were constructed by randomly sampling 1 Mbp blocks with replacement until the total sequence length was as close as possible to the size of the full genome (528,860,695 bp; Coffman *et al*. 2015). We also repeated the same procedure described above for choosing a step size for numerical differentiation (*ϵ* ∈ [10^−2^, …, 10^−7^] by factors of 10]). These bootstrapped frequency spectra were again used to conduct a likelihood ratio test between the models with and without inbreeding using the `LRT adjust()` method in ∂*a*∂*i* and comparing the test statistic to a weighted sum of *χ*^2^ distributions with zero and one degrees of freedom (see section above for rationale; Ota *et al*. 2000): 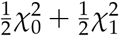.

#### Confidence Intervals for Composite Parameters

We used the constants listed above for sequence length (*L*), mutation rate (*µ*), and generation time (*g*) for pumas and domesticated cabbage to convert from the units used by ∂*a*∂*i* to real units of years and individuals. However, in order to estimate confidence intervals for these converted parameters, we need to correctly account for the fact that times and population sizes are products of two estimated parameters (either *θ* × *T*_∂*a*∂*i*_ or *θ* × *N*_∂*a*∂*i*_): 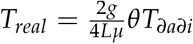and 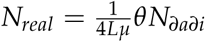. We do this by propagating the uncertainty in our estimates of each individual parameter into a combined estimate of the standard deviation for the composite parameter. In addition, because our original uncertainty estimates for each parameter were large and led to negative values in our confidence intervals, we instead estimated our uncertainty on a log scale. Taking the log of *T*_*real*_ and *N*_*real*_ gives us the following expressions for each parameter:

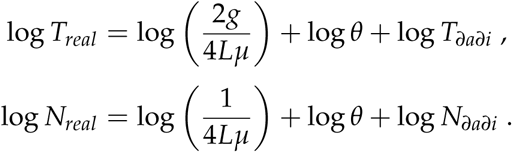

The corresponding expressions for the standard deviations of log *T*_*real*_ and log *N*_*real*_ are then

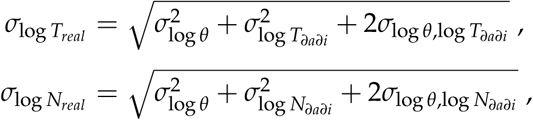

where 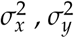, and *σ*_*x,y*_ are the variances and covariance for arbitrary variables *x* and *y*. With these new estimates of the standard deviation, we can obtain the log-scaled confidence intervals for *T*_*real*_ and 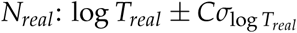 and 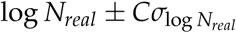. Here *C* is a constant chosen based on the desired confidence level (e.g., *C* = 1.96 for 95% confidence intervals). Exponentiating these expressions then gives us our confidence limits on the original scale.

## Data Availability

The inbreeding model is implemented in the Python package ∂*a*∂*i*, which is available on Bitbucket (https://bitbucket.org/gutenkunstlab/dadi). Code for generating and analyzing simulated and empirical data sets from this paper are available on GitHub (https://github.com/pblischak/inbreeding-sfs). The `bbc-shiny/` folder in the GitHub repository also contains a small R Shiny application for plotting the probability mass function of the *n*-fold convolution of beta-binomials with different sample sizes, allele frequencies, and inbreeding coefficients.

## Acknowledgements

The authors thank A. Ochoa, S. Turner-Hissong, M. Culver, O. Conejo, and J. Robinson for graciously agreeing to share empirical data for the present study. We would also like to thank members of the Barker and Gutenkunst labs for comments that helped to improve this manuscript. This work was supported by a National Science Foundation Postdoctoral Research Fellowship in Biology (IOS-1811784 to P.D.B.) and by the National Institute of General Medical Sciences of the National Institutes of Health (R01GM127348 to R.N.G.).

## Supplemental Tables

**Table S1:** Log-scale standard deviations for parameters in the model with inbreeding for American pumas across a series of step sizes.

| $\epsilon$ | $\sigma_{\log \nu_{TX}}$ | $\sigma_{\log \nu_{FL}}$ | $\sigma_{\log \tau_1}$ | $\sigma_{\log \tau_2}$ | $\sigma_{\log F_{TX}}$ | $\sigma_{\log F_{FL}}$ | $\sigma_{\log \theta}$ |
| --- | --- | --- | --- | --- | --- | --- | --- |
| $10^{-2}$ | 0.0604 | 1.281 | 0.1961 | 1.274 | 0.0385 | 0.0162 | 0.0061 |
| $10^{-3}$ | 0.0175 | 0.1738 | 0.0749 | 0.1993 | 0.0171 | 0.0427 | 0.0114 |
| $10^{-4}$ | 0.0104 | 0.3512 | 0.0209 | 0.3757 | 0.0277 | 0.0425 | 0.0109 |
| $10^{-5}$ | 0.0307 | 0.3501 | 0.0258 | 0.3741 | 0.0248 | 0.0425 | 0.0177 |
| $10^{-6}$ | 0.0595 | 0.3819 | 0.1853 | 0.4111 | 0.0330 | 0.0446 | 0.0671 |
| $10^{-7}$ | 0.0039 | 0.0086 | 0.0114 | 0.0052 | 0.0184 | 0.0222 | 0.0097 |

**Table S2:** Log-scale standard deviations for parameters in the model without inbreeding for American pumas across a series of step sizes.

| $\epsilon$ | $\sigma_{\log \nu_{TX}}$ | $\sigma_{\log \nu_{FL}}$ | $\sigma_{\log \tau_1}$ | $\sigma_{\log \tau_2}$ | $\sigma_{\log \theta}$ |
| --- | --- | --- | --- | --- | --- |
| $10^{-2}$ | 0.8452 | 1.057 | 1.894 | 1.067 | 0.1353 |
| $10^{-3}$ | 0.0642 | 0.0312 | 0.1588 | 0.0370 | 0.0160 |
| $10^{-4}$ | 0.1603 | 0.1758 | 0.3030 | 0.1818 | 0.0170 |
| $10^{-5}$ | 0.0205 | 0.0109 | 0.0410 | 0.0114 | 0.0105 |
| $10^{-6}$ | 0.0025 | 0.0079 | 0.0126 | 0.0029 | 0.0103 |
| $10^{-7}$ | 5.50e-5 | 6.69e-5 | 5.87e-5 | 1.03e-4 | 9.76e-3 |

**Table S3:** Log-scale standard deviations for parameters in the model with inbreeding for domesticated cabbage across a series of step sizes.

| $\epsilon$ | $\sigma_{\log \nu_1}$ | $\sigma_{\log \nu_2}$ | $\sigma_{\log \tau_1}$ | $\sigma_{\log \tau_2}$ | $\sigma_{\log F}$ | $\sigma_{\log \theta}$ |
| --- | --- | --- | --- | --- | --- | --- |
| $10^{-2}$ | 0.0432 | 1.913 | 0.1347 | 0.6388 | 0.0184 | 0.0171 |
| $10^{-3}$ | 0.0426 | 1.953 | 0.4052 | 0.8423 | 0.0187 | 0.0655 |
| $10^{-4}$ | 0.0434 | 1.411 | 0.2900 | 0.5930 | 0.0189 | 0.0470 |
| $10^{-5}$ | 0.0426 | 0.7802 | 0.2784 | 0.4184 | 0.0189 | 0.0453 |
| $10^{-6}$ | 0.0582 | 1.828 | 0.5530 | 0.1422 | 0.0183 | 0.0920 |
| $10^{-7}$ | 0.1664 | 0.0979 | 0.1425 | 0.4607 | 0.0147 | 0.0953 |

**Table S4:** Log-scale uncertainties for parameters in the model without inbreeding for domesticated cabbage across a series of step sizes.

| $\epsilon$ | $\sigma_{\log \nu_1}$ | $\sigma_{\log \nu_2}$ | $\sigma_{\log \tau_1}$ | $\sigma_{\log \tau_2}$ | $\sigma_{\log \theta}$ |
| --- | --- | --- | --- | --- | --- |
| $10^{-2}$ | 0.2236 | 0.0344 | 0.0597 | 0.0856 | 0.0161 |
| $10^{-3}$ | 2.068 | 4.006 | 0.5373 | 3.679 | 0.0512 |
| $10^{-4}$ | 2.596 | 1.325 | 0.5685 | 1.560 | 0.0438 |
| $10^{-5}$ | 3.435 | 10.57 | 0.7405 | 11.32 | 0.0496 |
| $10^{-6}$ | 0.9482 | 0.1776 | 0.2459 | 0.0701 | 0.0238 |
| $10^{-7}$ | 0.0071 | 0.0077 | 0.0191 | 0.0211 | 0.0179 |

## Supplemental Figures

**Figure S1:**
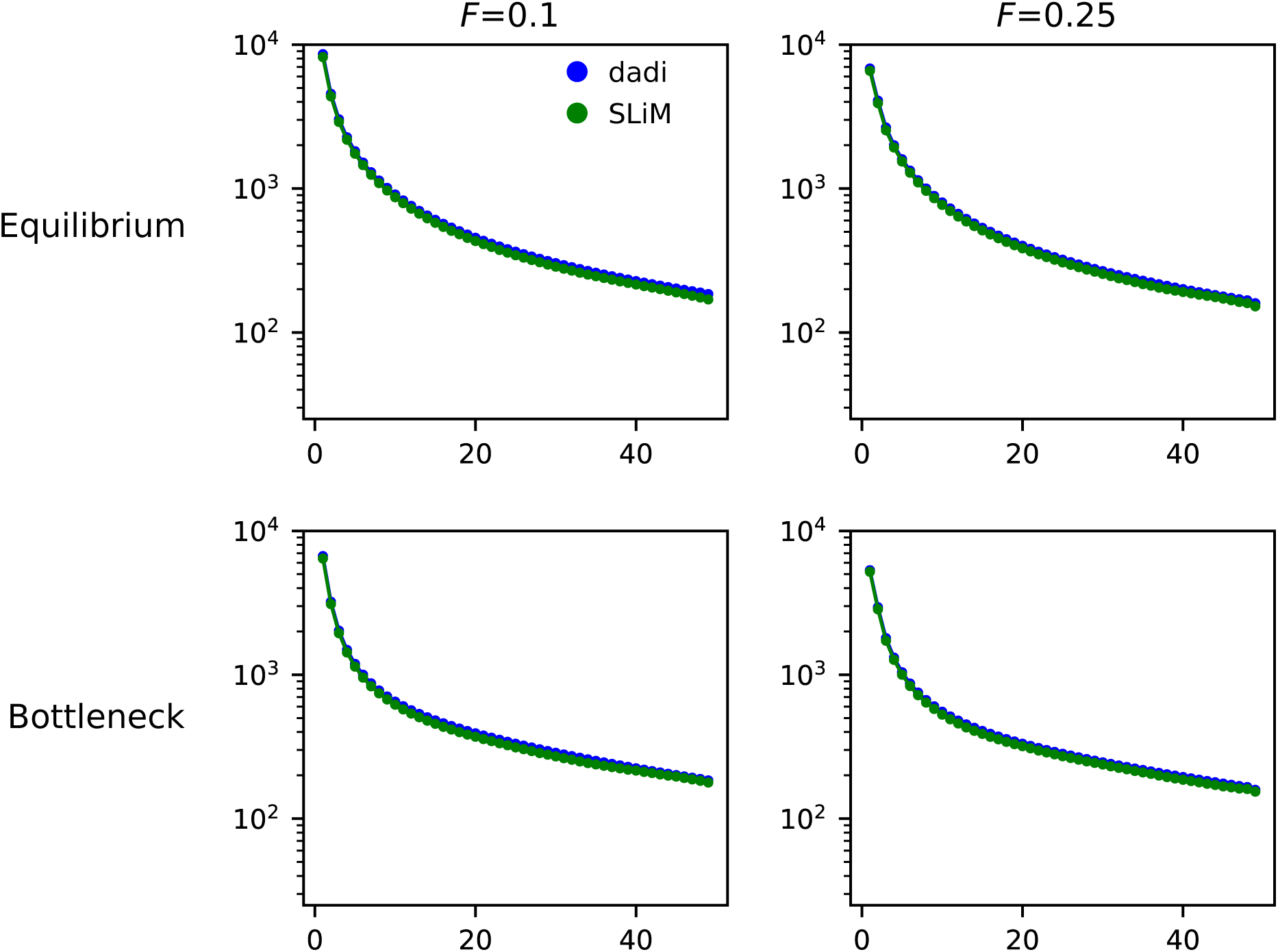
Comparison of expected frequency spectra for *F*=0.1 and 0.25 for ∂*a*∂*i* (blue) and SLiM (green) for the equilibrium and bottleneck+growth models.

**Figure S2:**
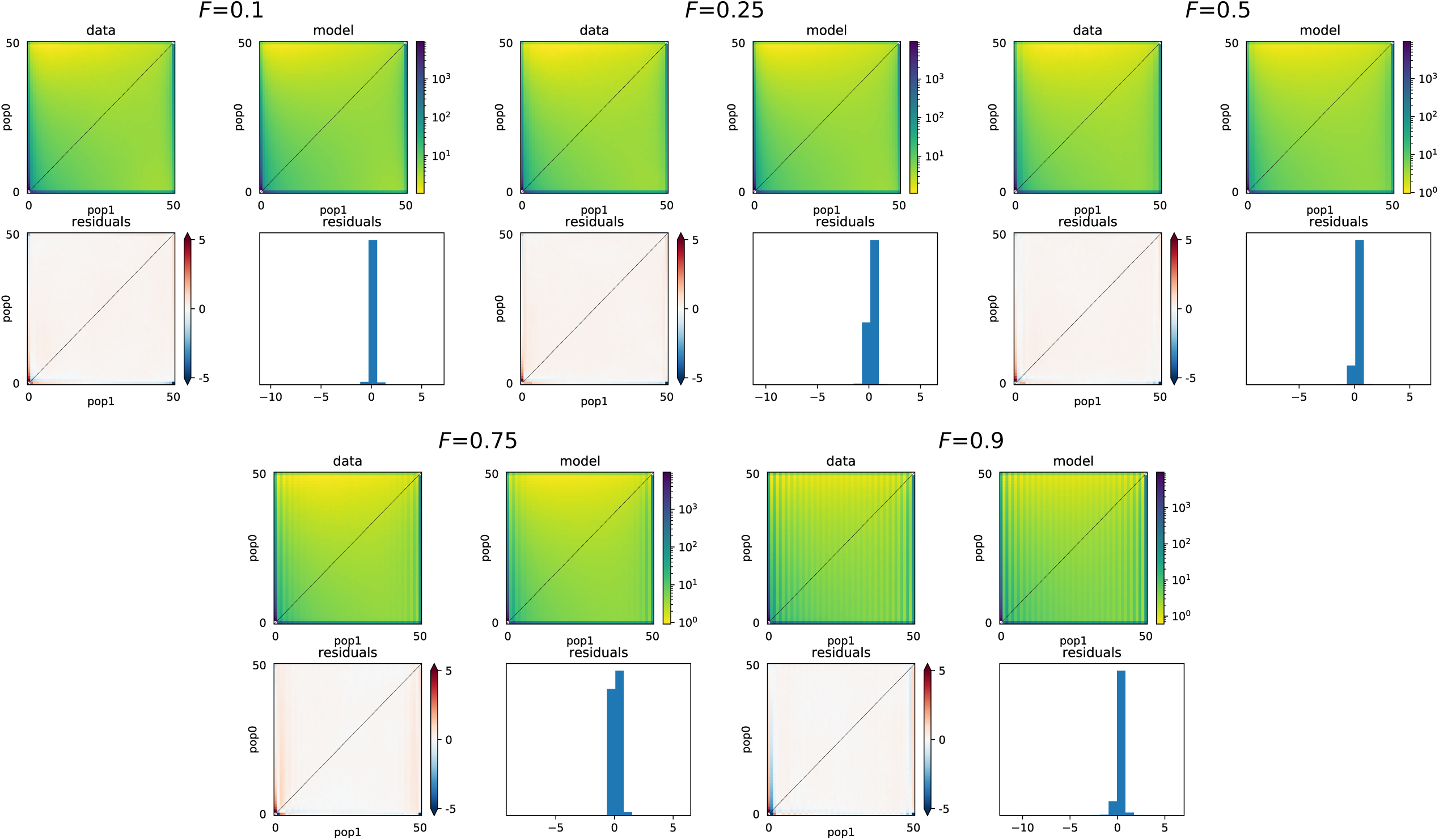
Comparison of expected frequency spectra for *F* = 0.1, 0.25, 0.5, 0.75, and 0.9 for ∂*a*∂*i* (‘model’) and SLiM (‘data’) for the domestication (divergence+one-way migration) model. Each plot also includes a heat map and histogram of the residual differences for the comparison between spectra from each method.

**Figure S3:**
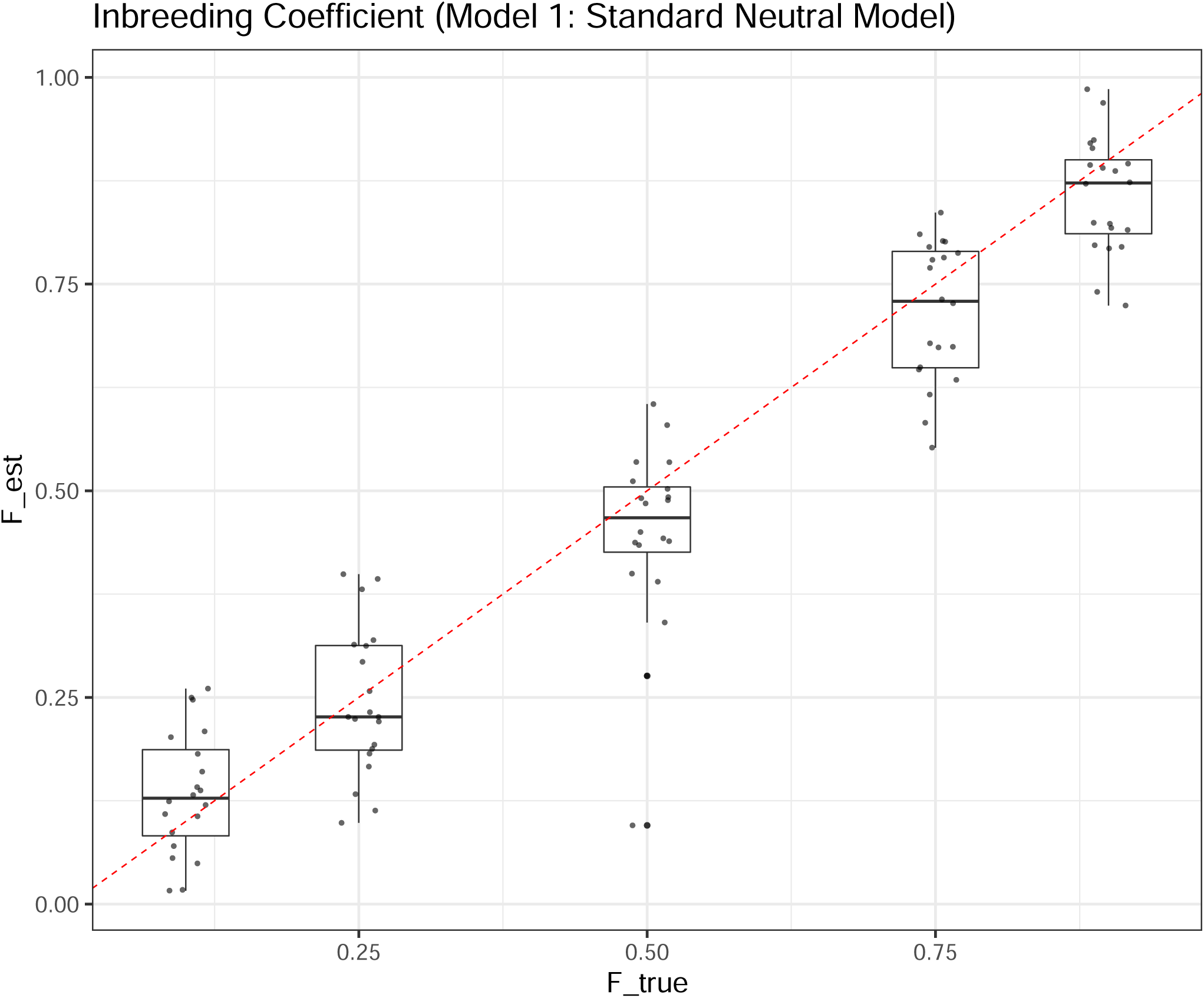
Estimates of the inbreeding coefficient for the equilibrium model generated by SLiM.

**Figure S4:**
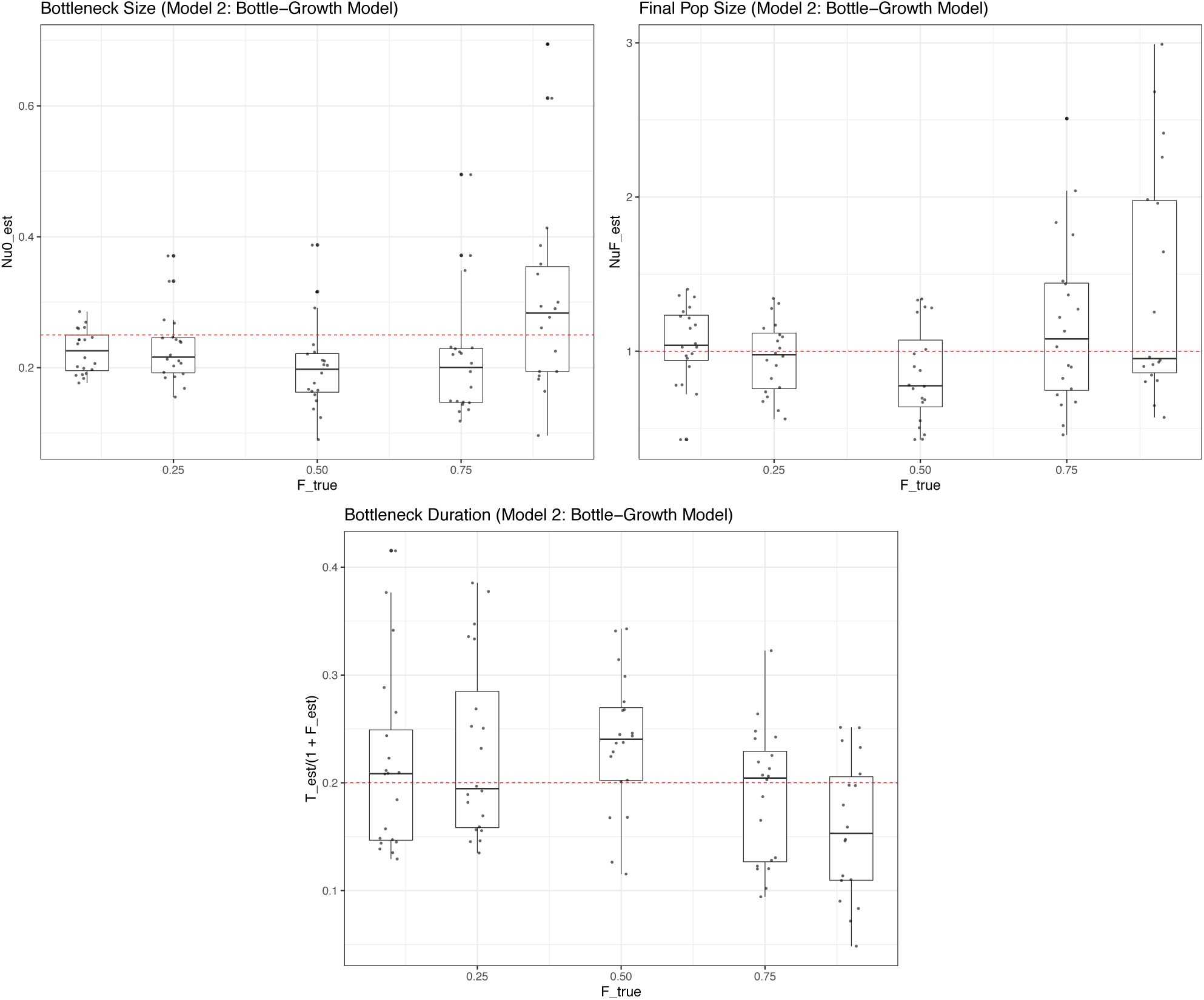
Parameter estimates from ∂*a*∂*i* for data simulated by SLiM for the bottleneck+growth model [bottleneck size (*ν*_0_): top left; final size (*ν*_*F*_): top right; scaled bottleneck duration 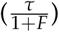: bottom].

**Figure S5:**
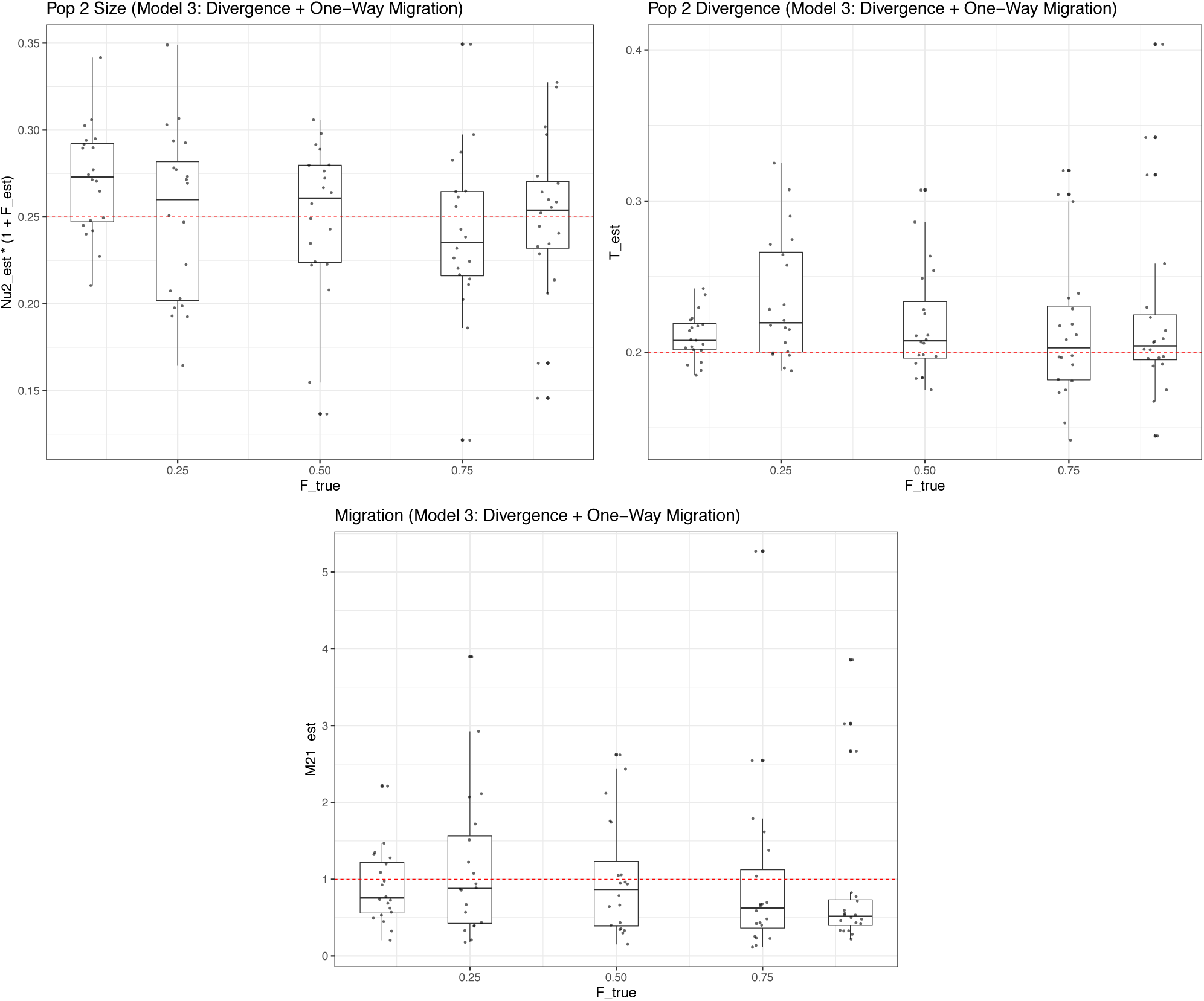
Parameter estimates from ∂*a*∂*i* for data simulated by SLiM for the domestication (divergence+one-way migration) model [scaled size of population two (*ν*_2_ × (1 + *F*)): top left; divergence time (*τ*): top right; migration (*M*_21_): bottom].

**Figure S6:**
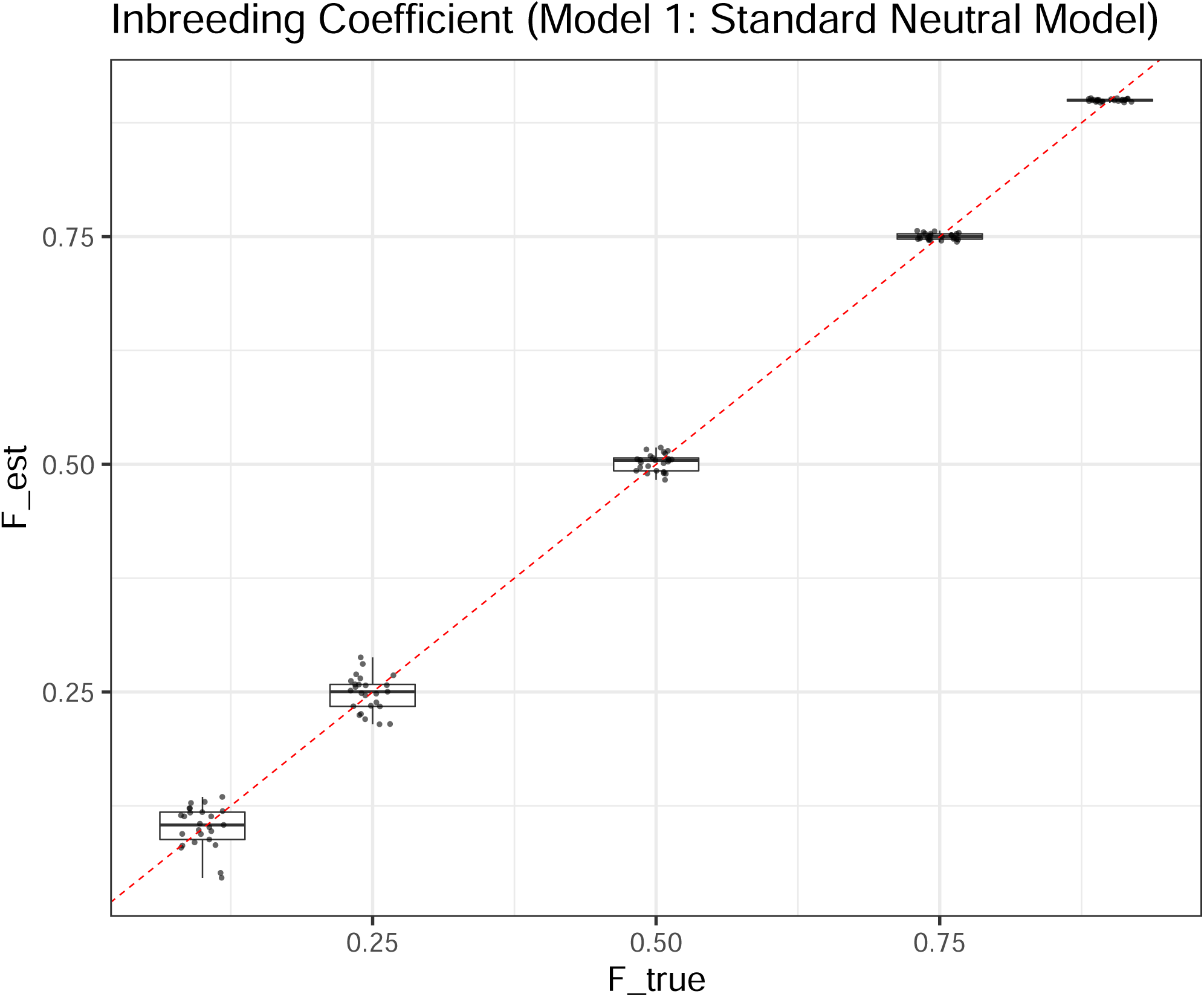
Estimates of the inbreeding coefficient for the standard neutral model for data generated by Poisson sampling within ∂*a*∂*i*.

**Figure S7:**
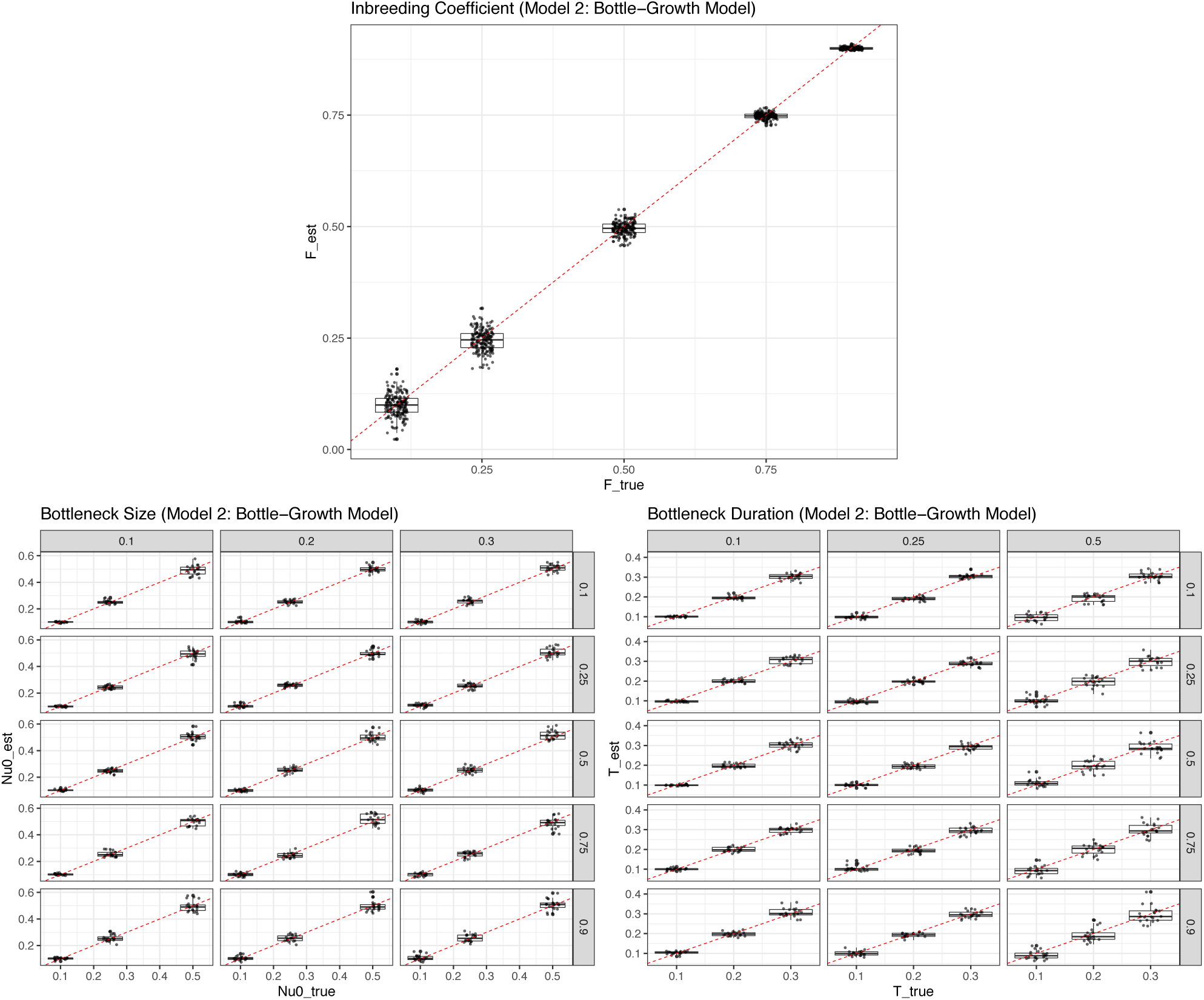
Parameter estimates from ∂*a*∂*i* for data generated by Poisson sampling for the bottleneck+growth model [inbreeding coefficient (*F*=0.1, 0.25, 0.5, 0.75, 0.9): top; bottleneck size (*ν*_0_=0.1, 0.25, 0.5): bottom left; bottleneck duration (*τ*=0.1, 0.2, 0.3): bottom right]. Plots for *ν*_0_ and *τ* are split into rows (true inbreeding coefficient) and column (true simulated parameter value).

**Figure S8:**
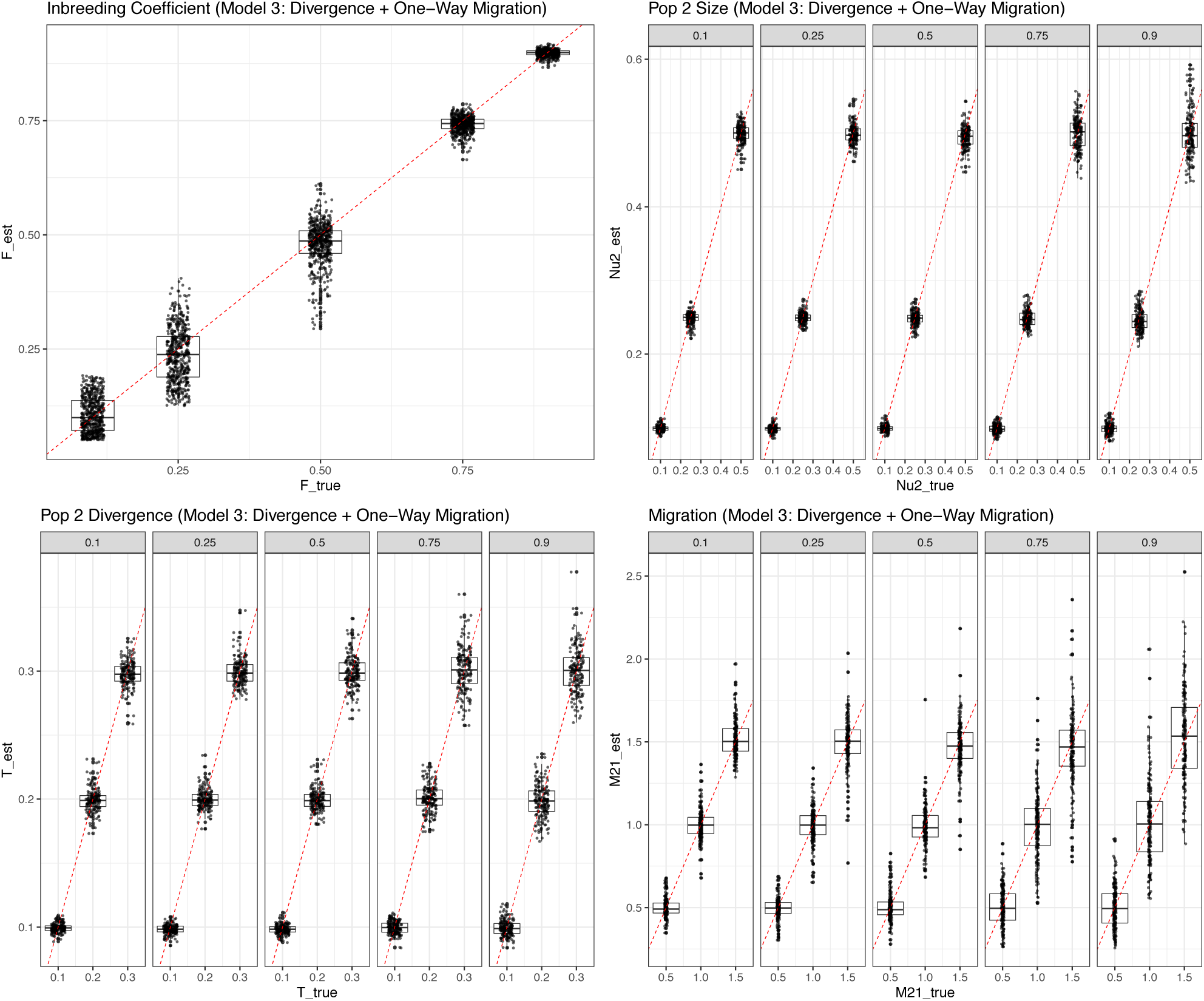
Parameter estimates from ∂*a*∂*i* for data generated by Poisson sampling for the divergence+one-way migration model [inbreeding coefficient (*F*=0.1, 0.25, 0.5, 0.75, 0.9): top left; size of population two (*ν*_2_=0.1, 0.25, 0.5): top right; divergence time (*τ*=0.1, 0.2, 0.3): bottom left; migration (*M*_21_ = 0.5, 1.0, 1.5): bottom right]. Plots for *ν*_2_, *τ*, and *M*_21_ are split into columns for each value of the inbreeding coefficient.

**Figure S9:**
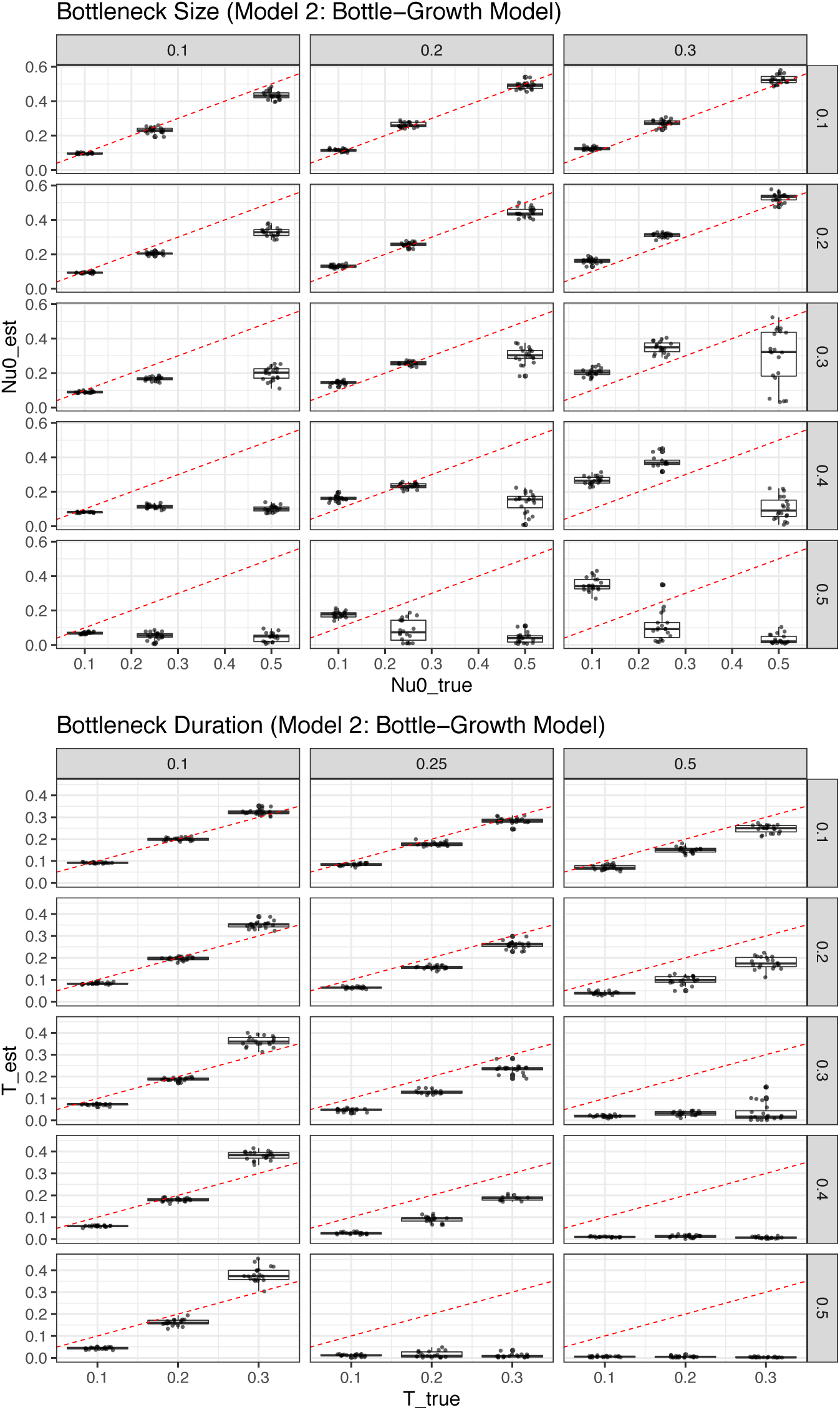
Parameter estimates from ∂*a*∂*i* for data generated by Poisson sampling for the bottleneck+growth model *when inbreeding is ignored* [inbreeding coefficient (*F*=0.1, 0.2, 0.3, 0.4, 0.5): *not estimated*; bottleneck size (*ν*_0_=0.1, 0.25, 0.5): top; bottleneck duration (*τ*=0.1, 0.2, 0.3): bottom]. Plots for *ν*_0_ and *τ* are split into rows (true inbreeding coefficient) and column (true simulated parameter value).

**Figure S10:**
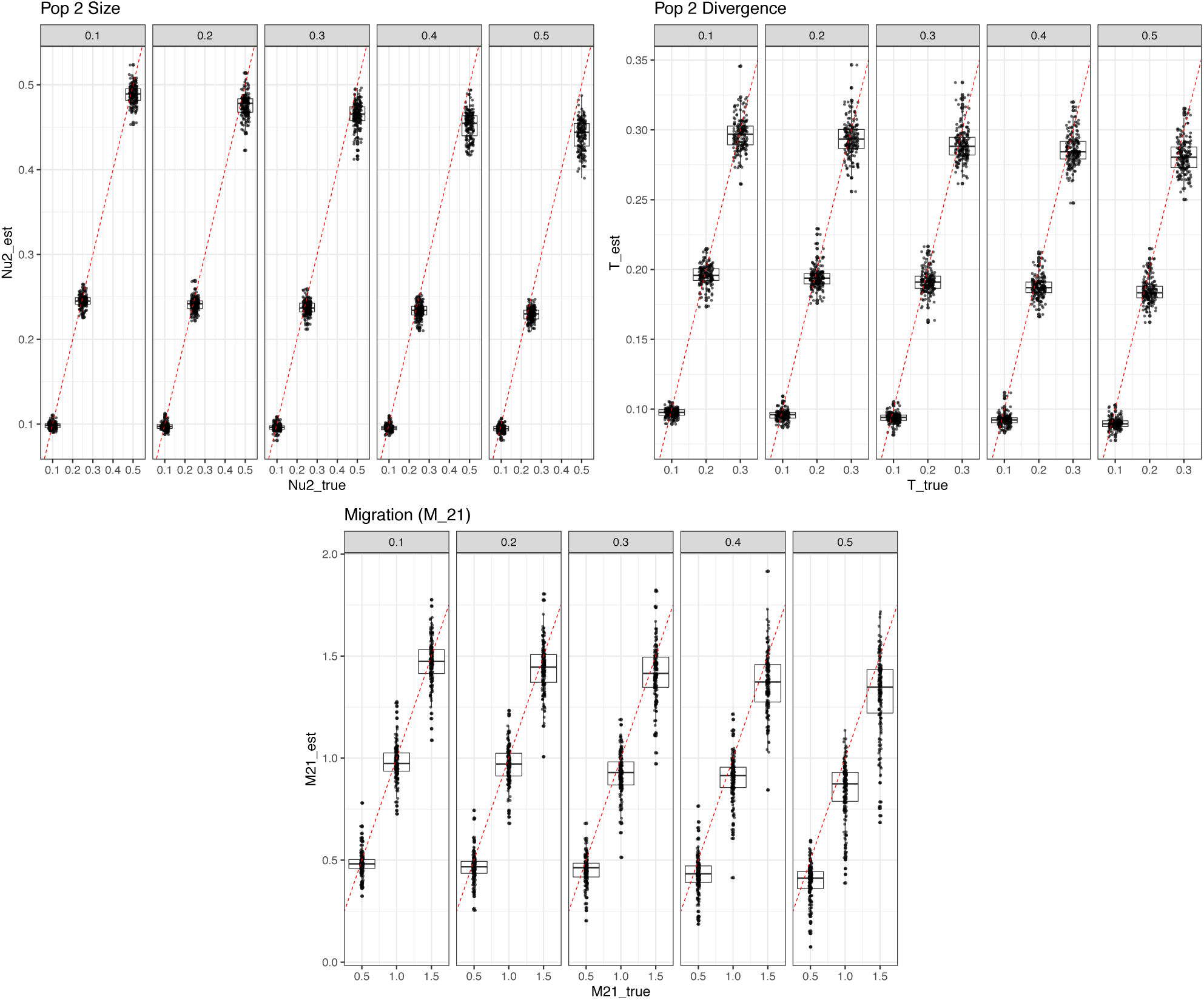
Parameter estimates from ∂*a*∂*i* for data generated by Poisson sampling for the divergence+one-way migration model *when inbreeding is ignored* [inbreeding coefficient (*F*=0.1, 0.2, 0.3, 0.4, 0.5): *not estimated*; size of population two (*ν*_2_=0.1, 0.25, 0.5): top left; divergence time (*τ*=0.1, 0.2, 0.3): top right; migration (*M*_21_ = 0.5, 1.0, 1.5): bottom]. Plots for *ν*_2_, *τ*, and *M*_21_ are split into columns for each value of the inbreeding coefficient.

**Figure S11:**
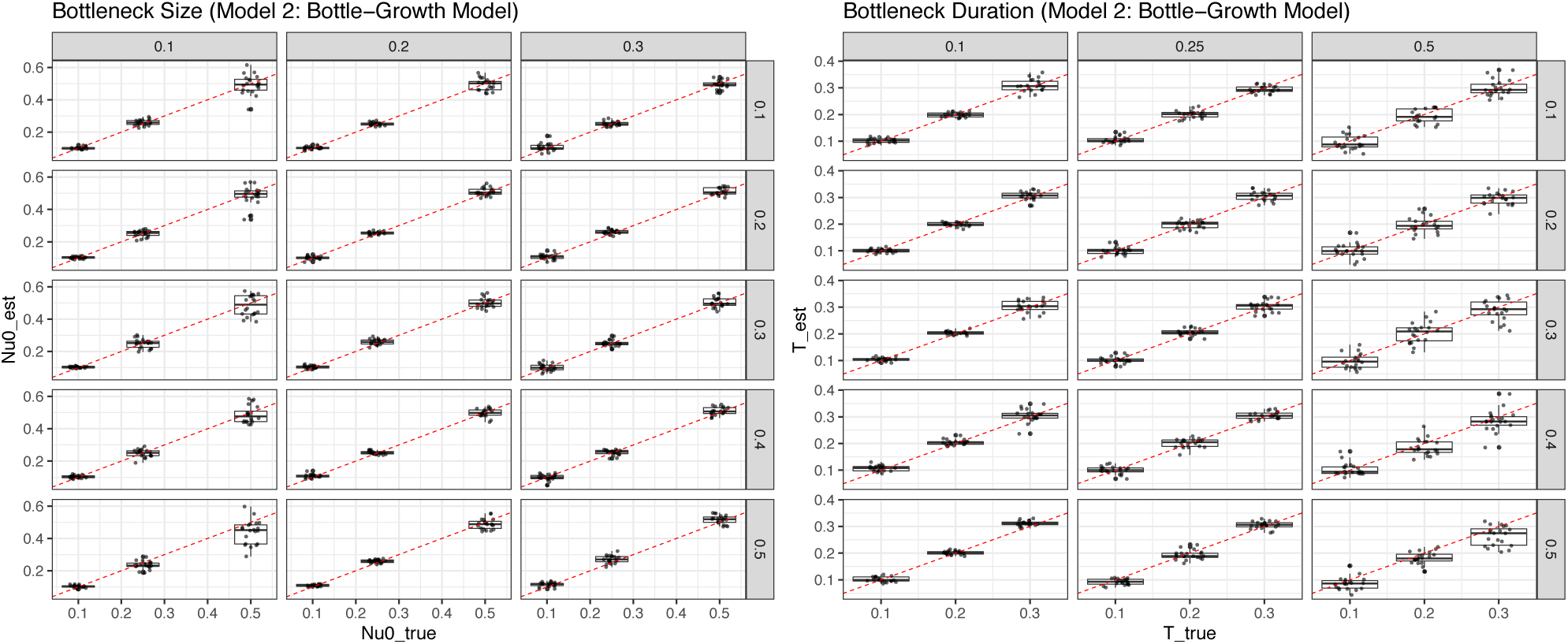
Parameter estimates from ∂*a*∂*i* for data generated by Poisson sampling for the bottle-neck+growth model *when rare variants are masked* [inbreeding coefficient (*F*=0.1, 0.2, 0.3, 0.4, 0.5): *not estimated*; bottleneck size (*ν*_0_=0.1, 0.25, 0.5): left; bottleneck duration (*τ*=0.1, 0.2, 0.3): right]. Plots for *ν*_0_ and *τ* are split into rows (true inbreeding coefficient) and column (true simulated parameter value).

**Figure S12:**
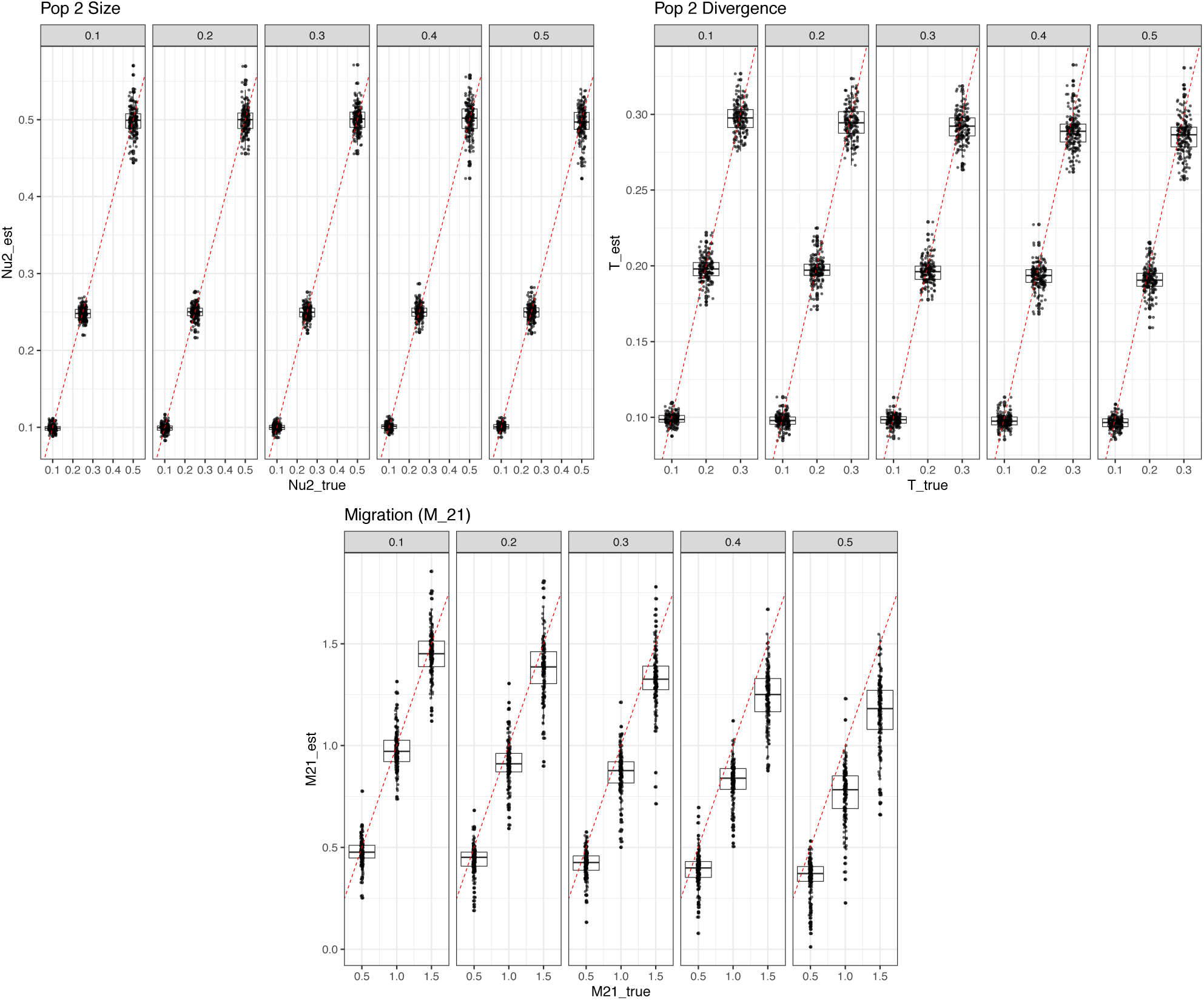
Parameter estimates from ∂*a*∂*i* for data generated by Poisson sampling for the divergence+one-way migration model *when rare variants are masked* [inbreeding coefficient (*F*=0.1, 0.2, 0.3, 0.4, 0.5): *not estimated*; size of population two (*ν*_2_=0.1, 0.25, 0.5): top left; divergence time (*τ*=0.1, 0.2, 0.3): top right; migration (*M*_21_ = 0.5, 1.0, 1.5): bottom]. Plots for *ν*_2_, *τ*, and *M*_21_ are split into columns for each value of the inbreeding coefficient.

**Figure S13:**
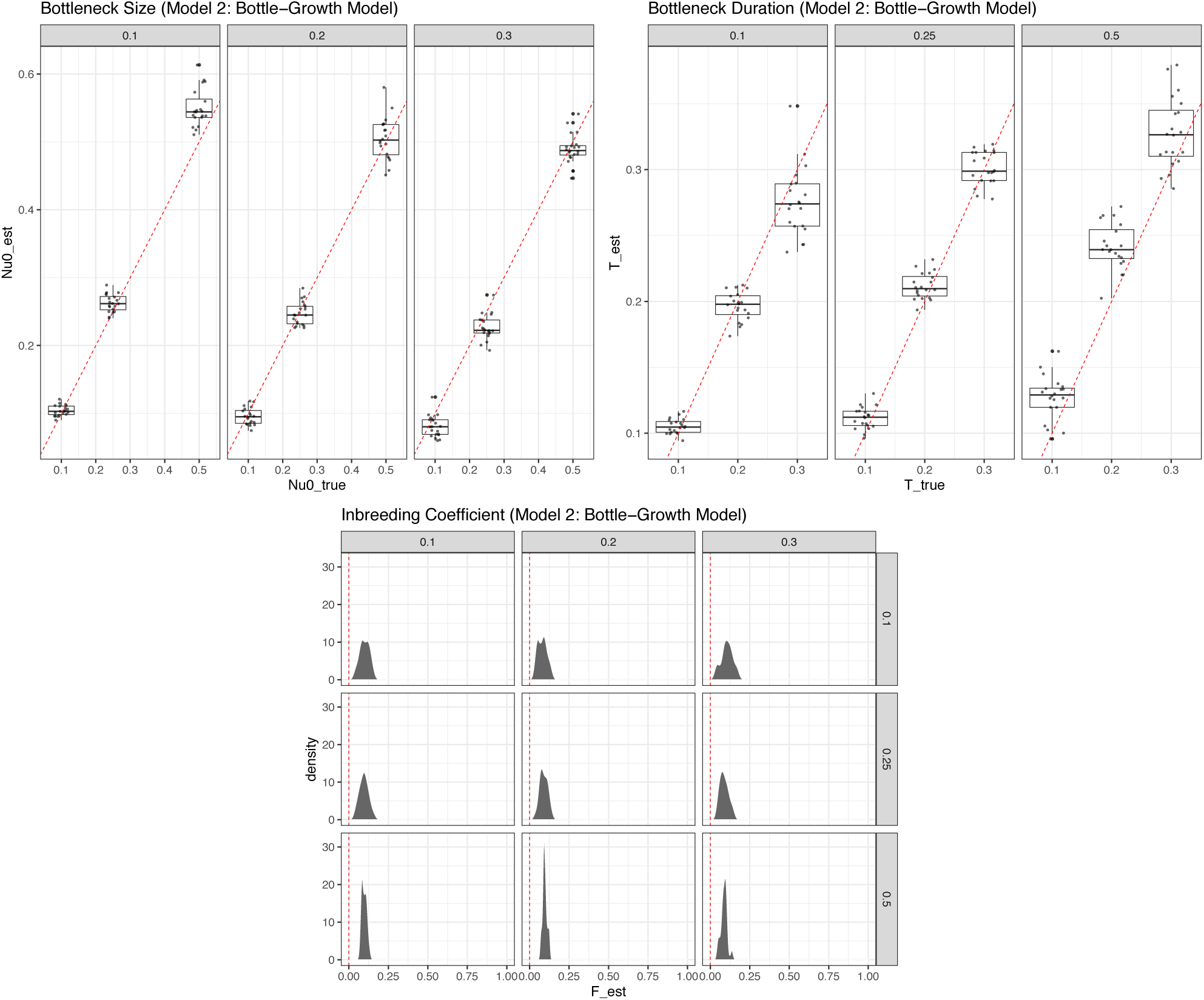
Parameter estimates from ∂*a*∂*i* for data generated by Poisson sampling for the bottle-neck+growth model *when inbreeding is absent but is still inferred* [bottleneck size (*ν*_0_=0.1, 0.25, 0.5): top left; bottleneck duration (*τ*=0.1, 0.2, 0.3): top right; inbreeding coefficient (*F*=0): bottom]. Plots for *ν*_0_ and *τ* are split into columns for each value of the other simulated parameter value.

**Figure S14:**
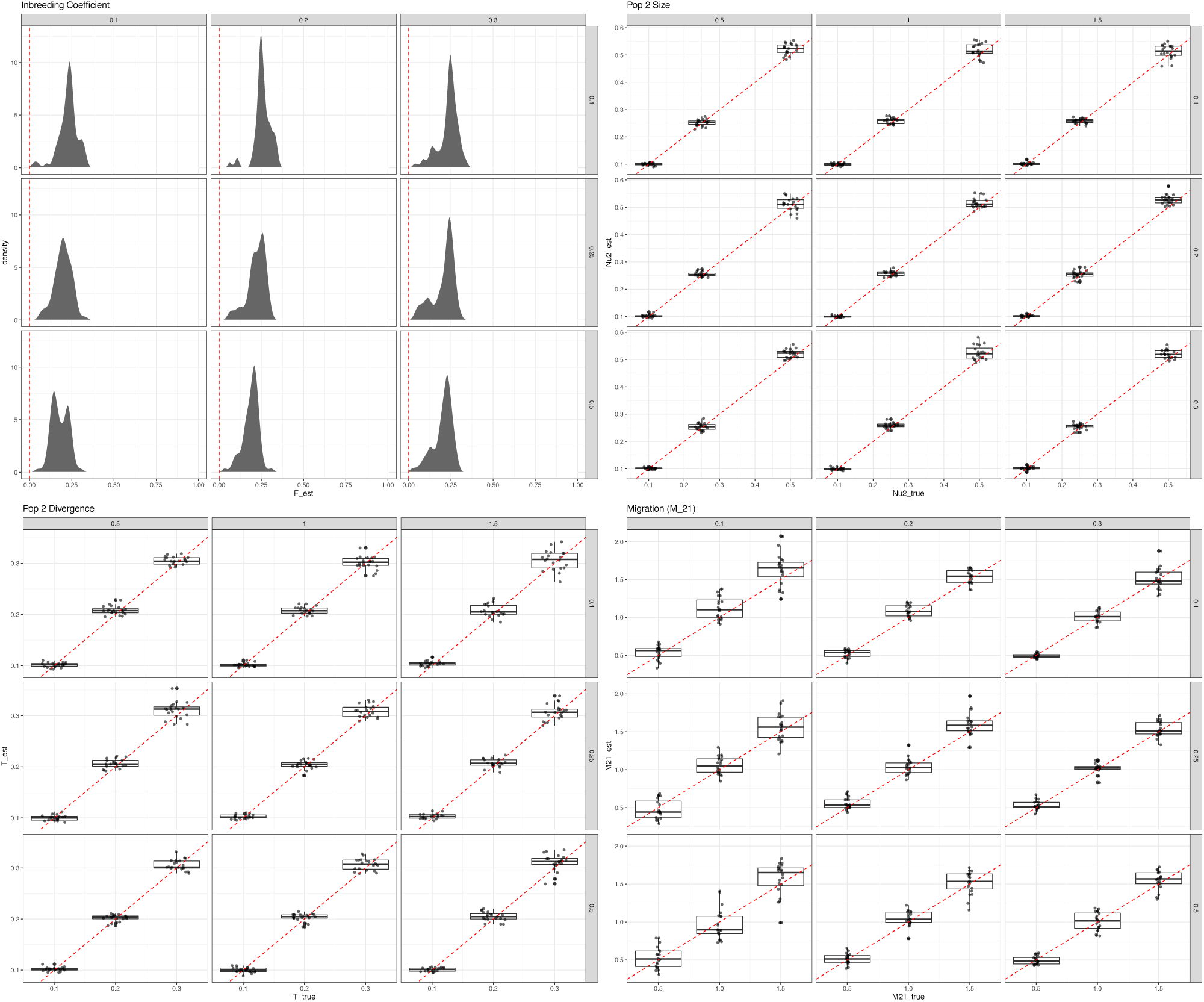
Parameter estimates from ∂*a*∂*i* for data generated by Poisson sampling for the divergence+one-way migration model *when inbreeding is absent but is still inferred* [inbreeding coefficient (*F*=0): top left; size of population two (*ν*_2_=0.1, 0.25, 0.5): top right; divergence time (*τ*=0.1, 0.2, 0.3): bottom left; migration (*M*_21_ = 0.5, 1.0, 1.5): bottom]. Plots for *ν*_2_, *τ*, and *M*_21_ are split into rows and columns for each value of the other two simulated parameters.

## References

Adams, A. M. and Hudson, R. R. 2004. Maximum-likelihood estimation of demographic parameters using the frequency spectrum of unlinked single-nucleotide polymorphisms. Genetics, 168: 1699–1712.

Balding, D. J. and Nichols, R. A. 1995. A method for quantifying differentiation between populations at multi-allelic loci and its implications for investigating identity and paternity. Genetica, 96: 3–12.

Balding, D. J. and Nichols, R. A. 1997. Significant genetic correlations among Caucasians at forensic DNA loci. Heredity, 108: 583–589.

Beissinger, T. M., Wang, L., Crosby, K., Durvasula, A., Hufford, M. B., and Ross-Ibarra, J. 2016. Recent demography drives changes in linked selection across the maize genome. Nature Plants, 2: 16084.

Belser, C., Istace, B., Denis, E., Dubarry, M., Baurens, F.-C., Falentin, C., Genete, M., Berrabah, W., Chèvre, A.-M., Delourme, R., Deniot, G., Denoeud, F., Duffé, P., Engelen, S., Lemainque, A., Manzanares-Dauleux, M., Martin, G., Morice, J., Noel, B., Vekemans, X., D’Hont, A., Rousseau-Gueutin, M., Barbe, V., Cruaud, C., Wincker, P., and Aury, J.-M. 2018. Chromosome-scale assemblies of plant genomes using nanopore long reads and optical maps. Nature Plants, 4: 879–887.

Browning, S. R., Browning, B. L., Daviglus, M. L., Daviglus, M. L., Durazo-Arvizu, R. A., Schneiderman, N., Kaplan, R. C., and Laurie, C. C. 2018. Ancestry-specific recent effective population size in the Americas. PLoS Genetics, 14: e1007385.

Caicedo, A. L., Williamson, S. H., Hernandez, R. D., Boyko, A., Fledel-Alon, A., York, T. L., Polato, N. R., Olsen, K. M., Nielsen, R., McCouch, S. R., Bustamante, C. D., and Purugganan, M. D. 2007. Genome-wide patterns of nucleotide polymorphism in domesticated rice. PLoS Genetics, 3: e163.

Ceballos, F. C., Joshi, P. K., Clark, D. W., Ramsay, M., and Wilson, J. F. 2018. Runs of homozygosity: Windows into population history and trait architecture. Nature Reviews Genetics, 19: 220–234.

Charlesworth, B. 1992. Evolutionary rates in partially self-fertilizing species. The American Naturalist, 140: 126–148.

Charlesworth, D. 2003. Effects of inbreeding on the genetic diversity of populations. Philosophical Transactions of the Royal Society B, 358: 1051–1070.

Cheng, F., Wu, J., Cai, C., Fu, L., Liang, J., Borm, T., Zhuang, M., Zhang, Y., Zhang, F., Bonnema, G., and Wang, X. 2016a. Genome resequencing and comparative variome analysis in a *Brassica rapa* and *Brassica oleracea* collection. Scientific Data, 3: 160119.

Cheng, F., Sun, R., Hou, X., Zheng, H., Zhang, F., Zhang, Y., Liu, B., Liang, J., Zhuang, M., Liu, Y., Liu, D., Wang, X., Li, P., Liu, Y., Lin, K., Bucher, J., Zhang, N., Wang, Y., Wang, H., Deng, J., Liao, Y., Wei, K., Zhang, X., Fu, L., Hu, Y., Liu, J., Cai, C., Zhang, S., Zhang, S., Li, F., Zhang, H., Zhang, J., Guo, N., Liu, Z., Liu, J., Sun, C., Ma, Y., Zhang, H., Cui, Y., Freeling, M. R., Borm, T., Bonnema, G., Wu, J., and Wang, X. 2016b. Subgenome parallel selection is associated with morphotype diversification and convergent crop domestication in *Brassica rapa* and *Brassica oleracea*. Nature Genetics, 48: 1218–1224.

Clark, P. U., Dyke, A. S., Shakun, J. D., Carlson, A. E., Clark, J., Wohlfarth, B., Mitrovica, J. X., Hostetler, S. W., and McCabe, A. M. 2009. The last glacial maximum. Science, 325: 710–714.

Coffman, A. J., Hsieh, P. H., Gravel, S., and Gutenkunst, R. N. 2015. Computationally efficient composite likelihood statistics for demographic inference. Molecular Biology and Evolution, 33: 591–593.

Cornejo, O. E., Yee, M.-C., Dominguez, V., Andrews, M., Sockell, A., Strandberg, E., Livingstone, III, D., Stack, C., Romero, A., Umaharan, P., Royaert, S., Tawari, N. R., Ng, P., Gutierrez, O., Phillips, W., Mockaitis, K., Bustamante, C. D., and Motamayor, J. C. 2018. Population genomic analyses of the chocolate tree, *Theobroma cacao* L., provide insights into its domestication process. Communications Biology, 1: 167.

Culver, M., Johnson, W. E., Pecon-Slattery, J., and O’Brien, S. J. 2000. Genomic ancestry of the American puma (*Puma concolor*). The Journal of Heredity, 91: 186–197.

Danecek, P., Auton, A., Abecasis, G., Albers, C. A., Banks, E., DePristo, M. A., Handsaker, R. E., Lunter, G., Marth, G. T., Sherry, S. T., McVean, G., Durbin, R., and 1000 Genomes Project Analysis Group 2011. The variant call format and VCFtools. Bioinformatics, 27: 2156–2158.

Doebley, J. F., Gaut, B. S., and Smith, B. S. 2006. The molecular genetics of crop domestication. Cell, 127: 1309–1321.

Excoffier, L., Dupanloup, I., Huerta-Sánchez, E., Sousa, V. C., and Foll, M. 2013. Robust demographic inference from genomic and SNP data. PLOS Genetics, 9: e1003905.

Fortier, A. L., Coffman, A. J., Struck, T. J., Irby, M. N., Burguete, J. E. L., Ragsdale, A. P., and Gutenkunst, R. N. 2019. DFEnitely different: Genome-wide characterization of differences in mutation fitness effects between populations. bioRxiv, doi: 10.1101/703918.

Gaut, B. S., Seymour, D. K., Liu, Q., and Zhou, Y. 2018. Demography and its effects on genomic variation in crop domestication. Nature Plants, 4: 512.

Gerbault, P., Allaby, R. G., Boivin, N., Rudzinski, A., Grimaldi, I. M., Pires, J. C., Climer Vigueira, C., Dobney, K., Gremillion, K. J., Barton, L., Arroyo-Kalin, M., Purugganan, M. D., Rubio de Casas, R., Bollongino, R., Burger, J., Fuller, D. Q., Bradley, D. G., Balding, D. J., Richerson, P. J., Gilbert, M. T. P., Larson, G., and Thomas, M. G. 2014. Storytelling and story testing in domestication. Proceedings of the National Academy of Sciences U.S.A., 111: 6159–6164.

Gutenkunst, R. N., Hernandez, R. D., Williamson, S. H., and Bustamante, C. D. 2009. Inferring the joint demographic history of multiple populations from multidimensional SNP frequency data. PLoS Genetics, 5: e1000695.

Haller, B. C. and Messer, P. W. 2019. SLiM 3: Forward genetic simulations beyond the Wright-Fisher model. Molecular Biology and Evolution, 36: 632–637.

Hansen, K. 1992. Cougar, the American lion. Flagstaff, AZ: Northland Publishing.

Hartfield, M. and Bataillon, T. 2019. Selective sweeps under dominance and inbreeding. bioRxiv, doi: https://doi.org/10.1101/318410.

Hartfield, M. and Glémin, S. 2016. Limits to adaptation in partially selfing species. Genetics, 203: 959–974.

Hughes, P. D. and Woodward, J. C. 2017. Quaternary glaciation in the Mediterranean mountains: A new synthesis. Geological Society, London, Special Publications, 433: 1–23.

Hughes, P. D., Woodward, J. C., and Gibbard, P. L. 2006. Quaternary glacial history of the Mediter-ranean mountains. Progress in Physical Geography: Earth and Environment, 30: 334–364.

Hunter, J. D. 2007. Matplotlib: A 2d graphics environment. Computing in Science & Engineering, 9: 90–95.

Johnson, S. G. 2014. The NLopt nonlinear-optimization package, http://github.com/stevengj/nlopt.

Johnson, W. E., Onorato, D. P., Roelke, M. E., Land, E. D., Cunningham, M., Belden, R. C., McBride, R., Jansen, D., Lotz, M., Shindle, D., Howard, J., Wildt, D. E., Penfold, L. M., Hostetler, J. A., Oli, M. K., and O’Brien, S. J. 2010. Genetic restoration of the Florida panther. Science, 329: 1641–1645.

Jouganous, J., Long, W., Ragsdale, A. P., and Gravel, S. 2017. Inferring the joint demographic history of multiple populations: Beyond the diffusion approximation. Genetics, 206: 1549–1567.

Kardos, M., Qvarnström, A., and Ellegren, H. 2017. Inferring individual inbreeding and demo-graphic history from segments of identity by descent in *Ficedula* flycatcher genome sequences. Genetics, 205: 1319–1334.

Kim, B. Y., Huber, C. D., and Lohmueller, K. E. 2017. Inference of the distribution of selection coefficients for new nonsynonymous mutations using large samples. Genetics, 206: 345–361.

Kirin, M., McQuillan, R., Franklin, C. S., Campbell, H., McKeigue, P. M., and F.Wilson, J. 2010. Genomic runs of homozygosity record population history and consanguinity. PLoS ONE, 5: e13996.

Koenig, D., Hagmann, J., Li, R., Bemm, F., Slotte, T., Neuffer, B., Wright, S. I., and Weigel, D. 2019. Long-term balancing selection drives evolution of immunity genes in *Capsella*. eLife, 8: e43606.

Lukić, S. and Hey, J. 2012. Demographic inference using spectral methods on SNP data, with an analysis of the human out-of-Africa expansion. Genetics, 192(2): 619–639.

Maggioni, L. 2015. Domestication of Brassica oleracea L. Ph.D. thesis, Swedish University of Agricultural Sciences.

Meyer, R. S. and Purugganan, M. D. 2013. Evolution of crop species: Genetics of domestication and diversification. Nature Reviews Genetics, 14: 840–852.

Nielsen, R., Melissa J. Hubisz, I. H., Torgerson, D., Andrés, A. M., Albrechtsen, A., Gutenkunst, R., Adams, M. D., Cargill, M., Boyko, A., Indap, A., Bustamante, C. D.,, and Clark, A. G. 2009. Darwinian and demographic forces affecting human protein coding genes. Genome Research, 19: 838–849.

Nordborg, M. 2000. Linkage disequilibrium, gene trees and selfing: An ancestral recombination graph with partial self-fertilization. Genetics, 154: 923–929.

Nordborg, M. and Donnelly, P. 1997. The coalescent process with selfing. Genetics, 146: 1185–1195.

Ochoa, A., Onorato, D. P., Fitak, R. R., Roelke-Parker, M. E., and Culver, M. 2017. Evolutionary and Functional Mitogenomics Associated With the Genetic Restoration of the Florida Panther. Journal of Heredity, 108: 449–455.

Ochoa, A., Onorato, D. P., Fitak, R. R., Roelke-Parker, M. E., and Culver, M. 2019. *De novo* assembly and annotation from parental and F_1_ puma genomes for the Florida panther genetic restoration program. G3: Genes|Genomes|Genetics, 9: 3531–3536.

Ota, R., Waddell, P. J., Hasegawa, M., Shimodaira, H., and Kishino, H. 2000. Appropriate Likelihood Ratio Tests and Marginal Distributions for Evolutionary Tree Models with Constraints on Parameters. Molecular Biology and Evolution, 17: 798–803.

Pollack, E. 1987. On the theory of partially inbreeding finite populations. I. partial selfing. Genetics, 117: 353–360.

Powell, M. J. D. 2009. The BOBYQA algorithm for bound constrained optimization without derivatives. Technical Report 2009/NA06, Department of Applied Mathematics and Theoretical Physics, Cambridge University.

Quinlan, A. R. and Hall, I. M. 2010. BEDTools: A flexible suite of utilities for comparing genomic features. Bioinformatics, 26: 841–842.

R Core Team 2019. R: A language and environment for statistical computing. R Foundation for Statistical Computing, Vienna, Austria.

Robinson, J. A., Vecchyo, D. O.-D., Fan, Z., Kim, B. Y., von Holdt, B. M., Marsden, C. D., Lohmueller, K. E., and Wayne, R. K. 2016. Genomic flatlining in the endangered island fox. Current Biology, 26: 1183–1189.

Robinson, J. A., Räikkönen, J., Vucetich, L. M., Vucetich, J. A., Peterson, R. O., Lohmueller, K. E., and Wayne, R. K. 2019. Genomic signatures of extensive inbreeding in isle royale wolves, a population on the threshold of extinction. Science Advances, 5: eaau0757.

Robinson, J. D., Coffman, A. J., Hickerson, M. J., and Gutenkunst, R. N. 2014. Sampling strategies for frequency spectrum-based population genomic inference. BMC Evolutionary Biology, 4: 254.

Sawyer, S. A. and Hartl, D. L. 1992. Population genetics of polymorphism and divergence. Genetics, 132: 1161–1176.

Seal, U. S. and Lacy, R. C. 1994. A plan for genetic restoration and management of the Florida panther (Felis concolor coryi): report to the Florida Game and Freshwater Fish Commission. Conservation Breeding Specialist Group, Apple Valley, MN.

Shafer, A. B., Wolf, J. B., Alves, P. C., Bergström, L., Bruford, M. W., Brännström, I., Colling, G., Dalén, L., Meester, L. D., Ekblom, R., Fawcett, K. D., Fior, S., Hajibabaei, M., Hill, J. A., Hoezel, A. R., Höglund, J., Jensen, E. L., Krause, J., Kristensen, T. N., Krützen, M., McKay, J. K., Norman, A. J., Ogden, R., Österling, E. M., Ouborg, N. J., Piccolo, J., Popović, D., Primmer, C. R., Reed, F. A., Roumet, M., Salmona, J., Schenekar, T., Schwartz, M. K., Segelbacher, G., Senn, H., Thaulow, J., Valtonen, M., Veale, A., Vergeer, P., Vijay, N., Vilà, C., Weissensteiner, M., Wennerström, L., Wheat, C. W., and Zieliński, P. 2015. Genomics and the challenging translation into conservation practice. Trends in Ecology & Evolution, 30: 78–87.

Tataru, P., Mollion, M., Glémin, S., and Bataillon, T. 2017. Inference of distribution of fitness effects and proportion of adaptive substitutions from polymorphism data. Genetics, 207: 1103–1119.

Wickham, H. 2009. ggplot2: Elegant graphics for data analysis. Springer, New York.

Wickham, H., Averick, M., Bryan, J., Chang, W., McGowan, L. D., François, R., Grolemund, G., Hayes, A., Henry, L., Hester, J., Kuhn, M., Pedersen, T. L., Miller, E., Bache, S. M., Müller, K., Ooms, J., Robinson, D., Seidel, D. P., Spinu, V., Takahashi, K., Vaughan, D., Wilke, C., Woo, K., and Yutani, H. 2019. Welcome to the tidyverse. Journal of Open Source Software, 4: 1686.

Williamson, S. H., Hernandez, R., Fledel-Alon, A., Zhu, L., Nielsen, R., and Bustamante, C. D. 2005. Simultaneous inference of selection and population growth from patterns of variation in the human genome. Proceedings of the National Academy of Sciences U.S.A., 102: 7882–7887.

Wright, S. 1951. The genetical structure of populations. Annals of Eugenics, 15: 323–354.

Xue, Y., Prado-Martinez, J., Sudmant, P. H., Narasimhan, V., Ayub, Q., Szpak, M., Frandsen, P., Chen, Y., Yngvadottir, B., Cooper, D. N., de Manuel, M., Hernandez-Rodriguez, J., Lobon, I., Siegismund, H. R., Pagani, L., Quail, M. A., Hvilsom, C., Mudakikwa, A., Eichler, E. E., Cranfield, M. R., Marques-Bonet, T., Tyler-Smith, C., and Scally, A. 2015. Mountain gorilla genomes reveal the impact of long-term population decline and inbreeding. Science, 348: 242–245.

